# A constraint-based framework to reconstruct interaction networks in microbial communities

**DOI:** 10.1101/2024.01.30.577913

**Authors:** Omid Oftadeh, Asli Sahin, Evangelia Vayena, Vassily Hatzimanikatis

## Abstract

Microbial communities live in diverse habitats and significantly impact our health and the environment. However, the principles that govern their formation and evolution remain poorly understood. A crucial step in studying microbial communities is to identify the potential metabolic interactions between the community members, such as competition for nutrients or cross-feeding. Due to the size and complexity of the metabolic network of each organism, there may be a variety of connections between each pair of organisms, which poses a challenge to unraveling the metabolic interactions. Here, we present ReMIND, a computational framework to reconstruct the interaction networks in microbial communities based on the metabolic capabilities of individual organisms. We applied ReMIND to a well-studied uranium-reducing community and the honeybee gut microbiome. Our results provide new perspectives on the evolutionary forces that shape these ecosystems and the trade-off between metabolite exchange and biomass yield. By enumerating alternative interaction networks, we systematically identified the most likely metabolites to be exchanged and highlighted metabolites that could mediate competitive interactions. We envision that ReMIND will help characterize the metabolic capacity of individual members and elucidate metabolic interactions in diverse communities, thus holds the potential to guide many applications in precision medicine and synthetic ecology.

## Introduction

Microbial communities inhabit diverse ecological environments and significantly impact our health and environment^1–5^. The presence of different species in a microbial community gives rise to emergent properties that individual microbes do not possess. These properties, often the result of complex physicochemical interactions, lead to a range of effects, including enhanced growth^6^, increased resilience against perturbations^7^, increased virulence^8,9^, and the development of new biochemical capabilities^10–14^. These interactions, primarily mediated by small molecules or metabolites, are key to understanding the collective behavior of microbial communities.

Despite recent advances in the characterization of microbial exometabolomes^15–17^, the metabolic interactions in microbial communities and the principles governing the formation and evolution of such interactions remain poorly understood. In this regard, computational tools can enable us to unravel the inherent complexity of microbial communities and improve our understanding of the system. Genome-scale metabolic models (GEMs) are mathematical representations of the biochemical reactions that can occur in an organism according to the functional annotation of the organism’s genome and experimental evidence^18^. These models have been used extensively for a wide range of applications, including the study of microorganisms^19^ and human diseases^20,21^, metabolic engineering^22,23^, and drug target discovery^24^. GEMs have also been used to study and simulate microbial communities^25–29^, facilitated by recent advances in automated reconstruction of GEMs^30–32^ and the increasing availability of manually curated models^33^.

Metabolic models provide valuable information about metabolism, including nutrient uptake and by-product secretion. Various computational methods have used this information to identify metabolic exchanges in microbial communities. Despite differences in assumptions and methodologies, these methods typically involve two steps. The first step identifies boundary metabolites, i.e., metabolites that are present in the habitat independently of the community. These metabolites may originate from the host, be naturally present in the habitat, or be introduced by the growth medium. Previous studies have defined these boundary metabolites by minimizing the number of nutrients required to maintain a physiology^26,28^ or by assuming that only those metabolites that promote maximum growth are externally available^29,34^. After identifying the boundary metabolites, the next step is to identify metabolic interactions. To this end, some studies minimize the number of exchanged metabolites to reduce species interdependence^26,28^, while others use flux variability analysis^29,34^ or graph-based methods^25^ to detect overlapping metabolites between species. These methods have been successfully applied to specify interactions in communities ranging from two to forty species. However, none of these methods consider trade-offs between metabolite exchange and biomass yield. In addition, minimizing species interdependence is not always consistent with the driving forces of the community^27,35^, suggesting the need for a more comprehensive framework incorporating a wide range of objective functions.

In this study, we present ReMIND (Reconstruction of Microbial Interaction Networks using Decomposed in silico minimal exchanges) to systematically reconstruct metabolic interactions in microbial communities based on the metabolic capabilities of community members. Our framework identifies the boundary metabolites without enforcing growth optimality, allowing it to account for ecological conditions where growth is not optimal. We first address the problem at the species level by generating a set of alternative substrate and by-product profiles stratified based on biomass yield. We then reconstruct a community model using these generated profiles. We optimize a chosen objective function, which may reflect evolutionary goals, engineering objectives, or consistency with available data. We applied our framework to a well-studied two-species uranium-reducing community^36,37^ to evaluate the evolutionary history of the community and propose strategies to improve the community function. We also applied our framework to a larger honeybee gut microbiome composed of seven members^38,39^. Our framework sheds light on the complex interplay between biomass yield and metabolite exchange and identifies key metabolites that lead to cooperation or competition among species. Furthermore, the framework can guide experimental studies to reveal the underlying mechanisms of natural microbiomes and provide new insights into the design of synthetic communities.

## Results

### A generalized workflow to reconstruct microbial interaction networks

We have developed a computational constraint-based framework for reconstructing metabolic interaction networks in microbial communities. Our framework takes as input (i) the genome-scale metabolic models of individual species and (ii) a list defining the available extracellular metabolites. These metabolites may be provided by the host, the environment, or the other species in the community. Such a list may be defined by experimental studies (e.g., exometabolomics) or generated context-specifically for the community under study by focusing on the key metabolites (Figure 1 upper left).

**Figure 1:**
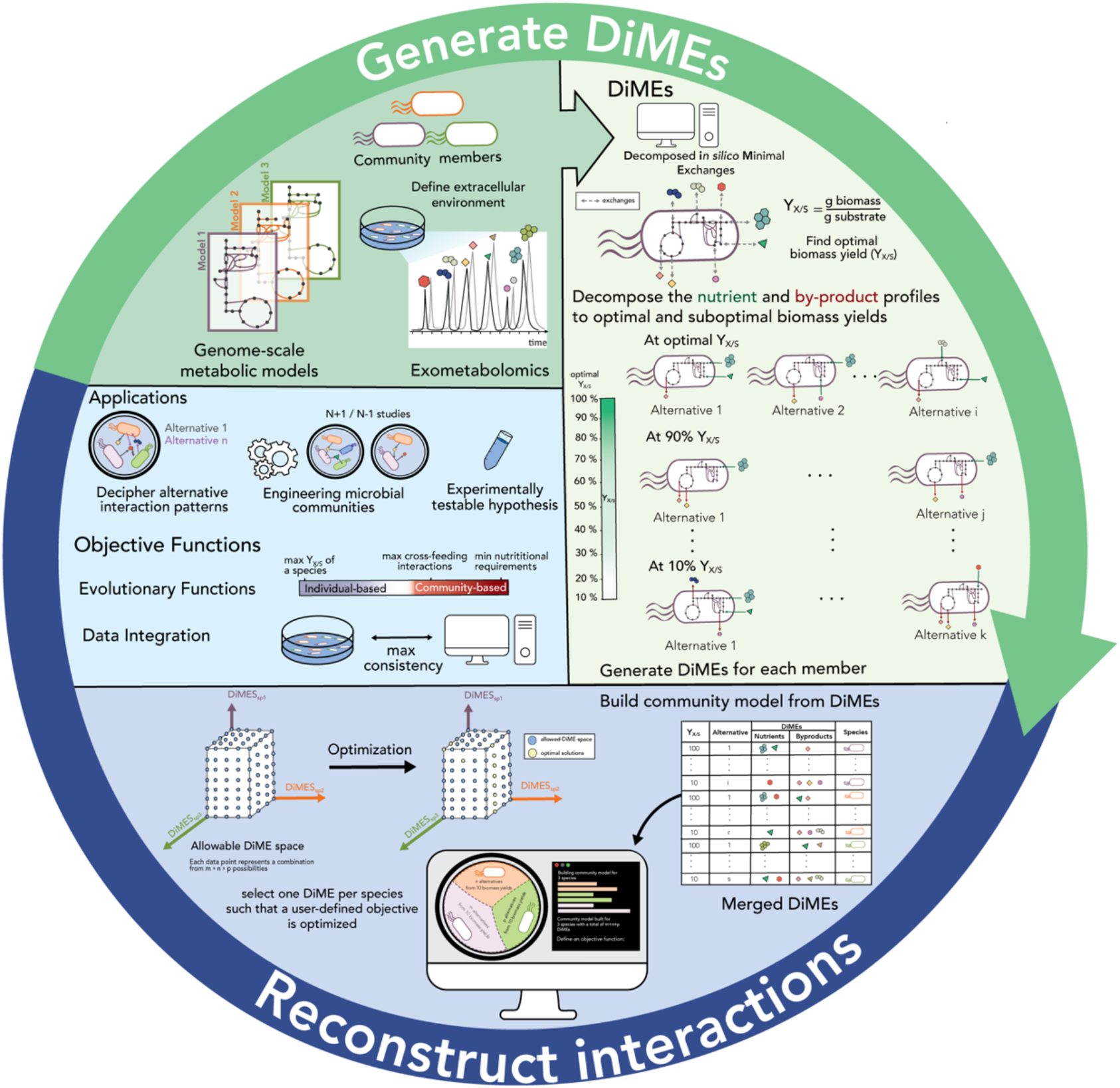
The workflow to reconstruct interaction networks in communities. From left in clockwise direction: GEMs of individual species and a list of available nutrients are the input to ReMIND. The ReMIND workflow starts with the generation of DiMEs. A community model is then built from the generated DiMEs. Following a user-defined objective function, metabolic interaction networks are reconstructed. ReMIND has diverse applications, from deciphering metabolic interaction networks, to engineering microbial communities with desired properties, and can propose experimentally testable hypothesis.

The framework starts with independently generating alternative nutrient and by-product profiles for each species. We aimed to create a framework not limited by the assumption of growth optimality for individual species within the community. To achieve this, we decomposed the nutrient and by-product profiles based on biomass yields. In this way, we account for optimal and sub-optimal yield regimes. Biomass yield can be calculated in different ways, e.g., based on all substrates, carbon, or nitrogen consumption, depending on the community under study. We named these substrate-byproduct profiles Decomposed in silico Metabolic Exchanges (DiMEs). The DiMEs represent all metabolic capabilities of a species in the given extracellular environment, with each DiME representing a specific physiology (Figure 1 upper right).

The next step is to assemble DiMEs into a community model. The DiMEs form the basis of the interaction space, where any combination of DiMEs of individual species creates an interaction profile. While there are many possible interactions, not all are biologically relevant. Therefore, we formulated an Integer-Linear Programming (ILP) problem to reconstruct relevant interaction networks. The ILP formulation selects one DiME per species to optimize an objective function. The objective function may represent a community driving force, a consistency score with partially observed interactions, or a desirable function.

### Specifying metabolic activity of a uranium-reducing community

We first used our method to reconstruct the interaction networks in a uranium-reducing community, including *G. sulfurreducens* and *R. ferrireducens*. These two species are significant components of a uranium-contaminated site in Rifle, Colorado^40^. The importance of *G. sulfurreducens* is due to its ability to reduce U (VI) to U (IV). *R. ferrireducens* cannot reduce uranium but competes with *G. sulfurreducens* over the shared nutrients. Previous studies reported that *G. sulfurreducens* and *R. ferrireducens* compete over acetate, ammonium, and Fe (III)^36^ (Figure 2a). Previous modeling efforts of this community assumed the two species interact only by competing over the abovementioned nutrients^37,41^. We used ReMIND to generate the DiMEs for *G. sulfurreducens* and *R. ferrireducens* (Table S1). We then studied the interaction networks that emerged, optimizing different objective functions.

**Figure 2:**
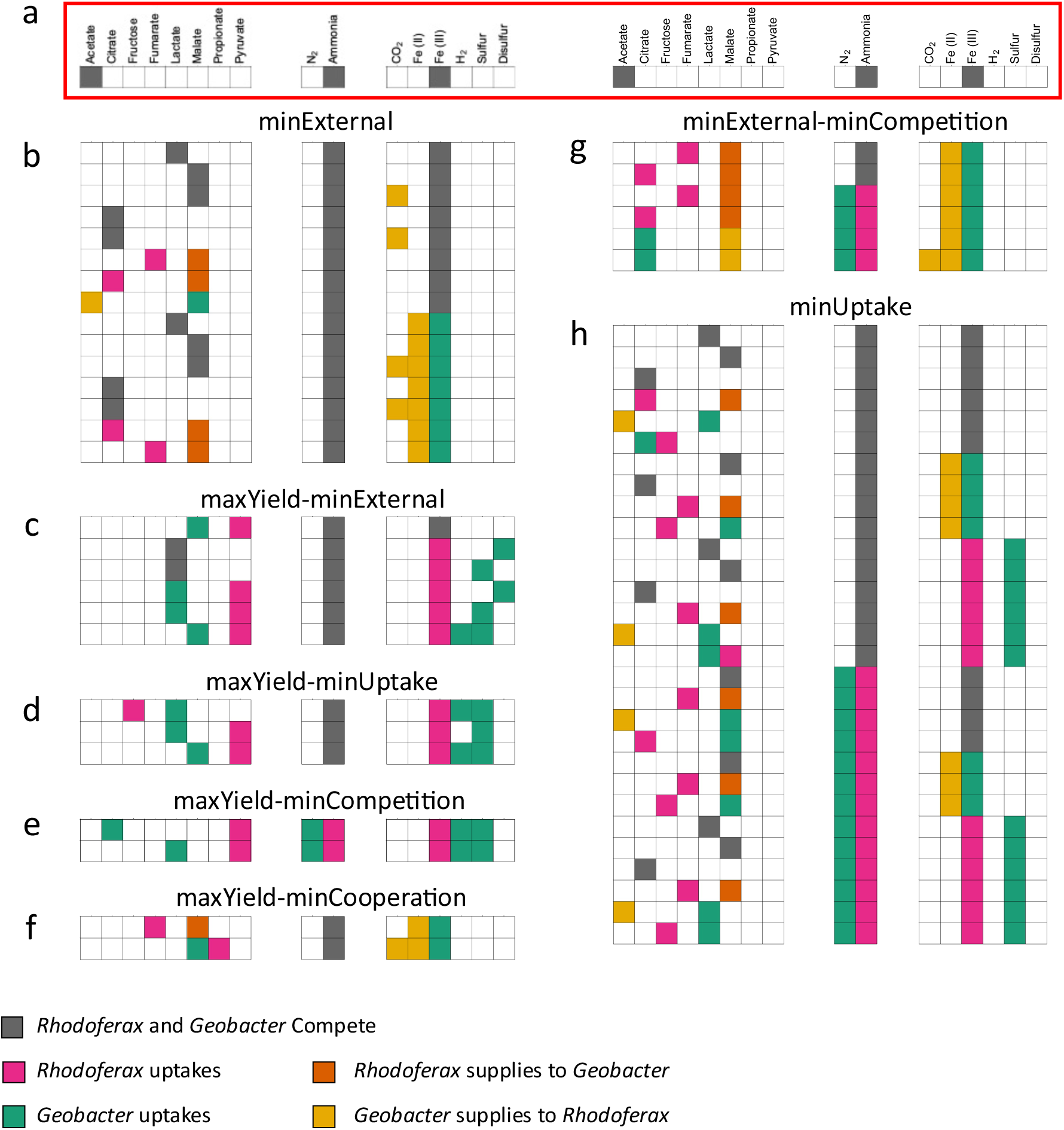
Illustration of interaction networks in the uranium-reducing community using Evolutionary objective functions. **a** The observed interactions between *G. sulfurreducens* and *R. ferrireducens,* which includes competition over acetate, ammonium, and Fe (III). The optimal solutions for different objective functions: **b** minimizing the number of external nutrients (minExternal), **c** maximizing the yield of both organisms while minimizing the number of external uptakes (maxYield-minExternal), **d** maximizing the yield of both organisms while minimizing the number of uptakes (maxYield-minUptake), **e** maximizing the yield of both organisms while minimizing the number of competitive interactions (maxYield-minCompetition), **f** maximizing the yield of both organisms while maximizing the number of cross-feeding interactions (maxYield-maxCrossfed), **g** minimizing the number of external nutrients and competitive interactions (minExternal-minCompetition), and **h** minimizing the number of uptaken nutrients (minUptake). Regarding the competition over ammonium, we observed three different patterns: (i) competition over ammonium in all optimal solutions (b, c, d, and f), (ii) switching between competition over ammonium and using different nitrogen sources, where *G. sulfurreducens* consumed N2 as an alternative nitrogen source to avoid competition (g and h), and (iii) no competition over ammonium (e). Regarding the competition over Fe (III), we observed two patterns: (i) competition over Fe (III) in some optimal solutions (b and c) and (ii) no competition over Fe (III) as the electron acceptor (d, e, f, g, and h). Interestingly, *G. sulfurreducens* and *R. ferrireducens* cooperated over iron in some cases, where *G. sulfurreducens* consumed Fe (III) and secreted Fe (II) which is then uptaken by *R. ferrireducens*. In such cases, *R. ferrireducens* relied on other electron acceptors. In all cases, we did not observe competition over acetate, reflecting that competition over acetate probably appeared due to the constrained availability of resources rather than optimality.

We generated DiMEs for *G. sulfurreducens* and *R. ferrireducens* for ten biomass yield regimes (Table S1). The number of unique DiMEs for *G. sulfurreducens* was 519, including 18 substrates and 16 by-products, where 8 metabolites could serve either as substrates or products in different alternatives. The number of unique DiMEs for *R. ferrireducens* was 400, including 19 substrates and 15 by-products, where 10 metabolites were common between substrates and products.

### Evolutionary objective functions can capture the uranium-reducing community driving force

We used the ILP formulation to perform three studies for this community using different sets of objective functions. First, we tried to capture the driving force of the community that led to the observed interactions. To this end, we defined seven evolutionary objective functions and generated alternative optimal solutions for each objective (Table *1*). If we could reproduce the observed interactions using an objective function, we could assume that this community had potentially evolved to optimize that objective function.

**Table 1:**
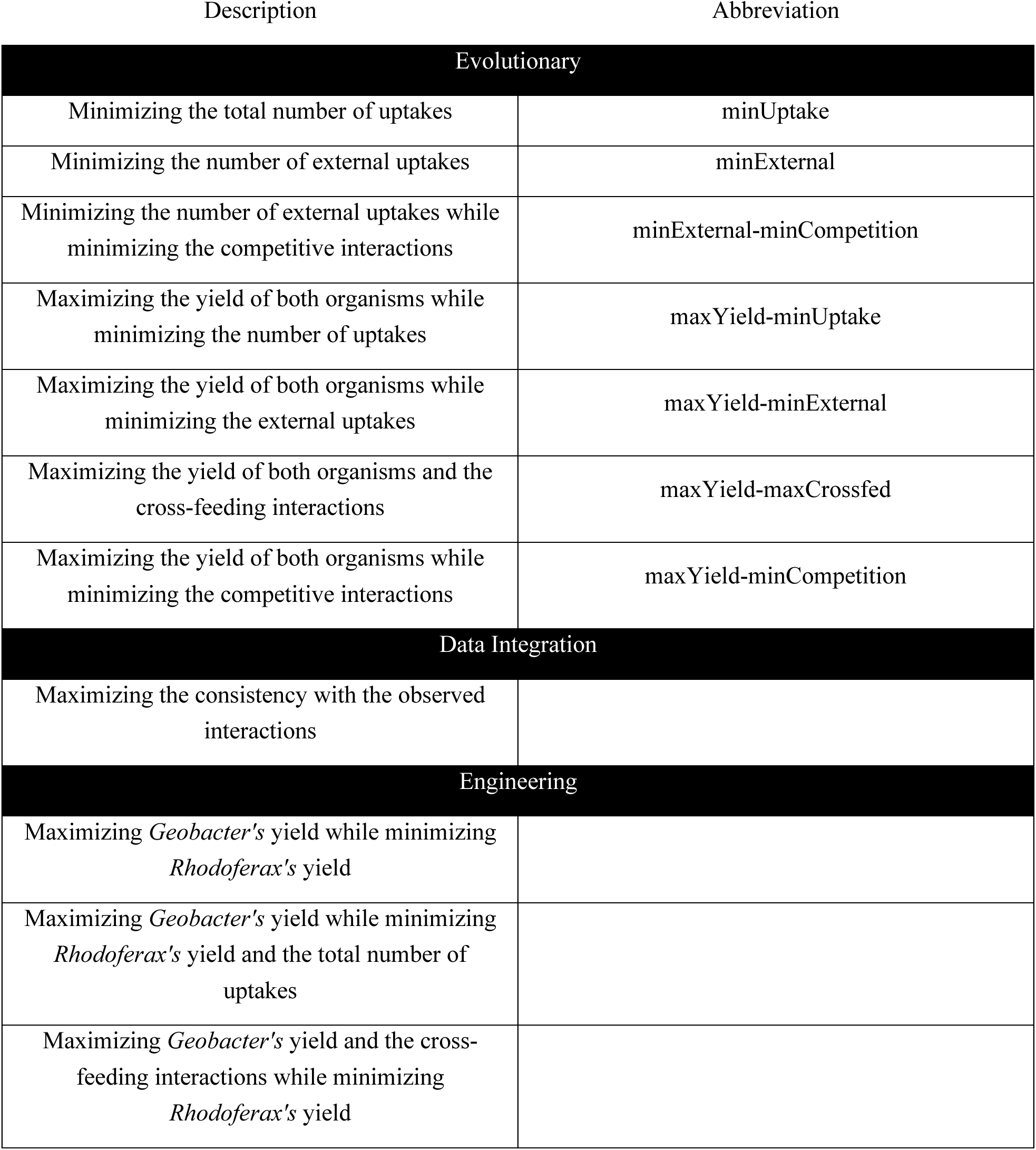
Different objective functions used with the ILP formulation to reconstruct the interaction network in the uranium-reducing community.

The first objective function was to minimize the sum of nutrient uptakes by individual species, denoted as minUptake, assuming that individual organisms evolved independently to minimize the resources they uptake. The second objective function was to minimize the uptake of external resources (abiotic) by the community (minExternal), assuming the two species evolved together to minimize the dependence on the environment. In addition to minimizing the uptake of external resources, the third objective minimizes the competitive interactions (minExternal-minCompetition), assuming further interdependence between the community members. The other four objective functions include maximizing the active yield regimes. They include (i) maximizing the yield regimes while minimizing the number of uptaken nutrients (maxYield-minUptake), (ii) maximizing the yield regimes while minimizing the number of external nutrients (maxYield-minExternal), (iii) maximizing the yield regimes while maximizing the cross-feeding interactions (maxYield-maxCrossfed), and (iv) maximizing the yield regimes while minimizing the competitive interactions (maxYield-minCompetition).

We generated the alternative optimal solutions for each objective function (Figure 2). Minimizing the number of external nutrients resulted in more competitive and cross-feeding interactions between the two species (Figure 2b, g) to reduce the community’s reliance on the environment. The cross-feeding interactions were mutual, indicating that *G. sulfurreducens* and *R. ferrireducens* can benefit from each other in an interdependent community. On the other hand, maximization of the yields indirectly reduced the cross-feeding interactions by driving the community to more selfish behavior (Figure 2c, d, e, f).

Three different patterns emerged concerning the competition over ammonium. The first pattern included an essential competition over ammonium, i.e., the two species competed over ammonium in all alternative optimal solutions. Four objective functions, minExternal, maxYield-minExternal, maxYield-minUptake, and maxYield-maxCrossfed, yielded the first pattern (Figure 2b, c, d, f). The second pattern featured alternative nitrogen sources for *G. sulfurreducens*. We then sought to explore the trade-off between the number of external sources and competitive interactions (i.e., minExternal-minCompetition). *G. sulfurreducens* could either compete with *R. ferrireducens* over ammonium or avoid competition by using N_2_ as the nitrogen source at the expense of increasing the number of external sources (Figure 2g). We observed the same pattern for minUptake, where we minimized the number of uptakes for each organism regardless of the interactions (Figure 2h). Thus, we can deduce that *G. sulfurreducens* could achieve the minimum number of substrates using either of the nitrogen sources. The third pattern was obtained when the objective function was maxYield-minCompetition, where no competition over ammonium was observed in the optimal solutions (Figure 2e). We also observed three patterns regarding the competition over Fe (III). The first pattern featured an essential cross-feeding interaction, where *G. sulfurreducens* consumed Fe (III) and secreted Fe (II), which was, in turn, consumed by *R. ferrireducens*. This pattern was observed for maxYield-maxCrossfed and minExternal-minCompetition (Figure 2f, g). In the second pattern, we observed no interactions over iron when the objective was maxYield-minUptake or maxYield-minCompetition (Figure 2d, e). The third pattern captured the competition over Fe (III) in some alternative solutions, while we observed either cooperation over Fe (II) or no interactions over iron in other alternatives. Three objective functions, minExternal, maxYield-minExternal, and minUptake, resulted in the third pattern. Finally, we did not observe the competition over acetate in the optimal solutions of these objective functions. This implies that competition over acetate is not an optimal phenotype, at least for these objective functions, and is probably observed due to the limited availability of other carbon sources in the environment.

Among the objective functions we considered, minExternal and maxYield-minExternal could capture the observed interactions better than the others (Figure 2b, c). These two objectives captured competition over ammonium (all alternatives) and Fe (III) (some alternatives), and both included minimizing the number of external nutrients. Given that *G. sulfurreducens* and *R. ferrireducens* minimized the number of uptaken external nutrients, we can conclude that the two species evolved in the same environment where the availability of nutrients was constrained.

### Inferring the metabolic landscape by integrating partial observations

In many cases, we have partial information about the interactions between community members. The systematic generation of DiMEs enables us to specify all potential interactions compatible with the partial information. To this end, we can formulate an objective function to find the most consistent interaction(s) with the observations (Table 1).

In the second study, we solved the ILP formulation, where we tried to find the most consistent interaction networks with the observations (see Methods). This problem had only one optimal solution, in which, in addition to the observed interactions, the two species also compete over sulfate and phosphate. We used the ILP formulation to find the DiMEs that support this set of interactions without establishing interactions through other metabolites. This included thirteen DiMEs for *G. sulfurreducens* and four DiMEs for *R. ferrireducens* (Figure 3a). The DiMEs for *G. sulfurreducens* and *R. ferrireducens* respectively corresponded to a range of suboptimal yield regimes from 10% to 60% and 10% to 40% of the optimal yield, which supports our finding in the previous section that the observed interactions were established mainly due to the constrained availability of the nutrients rather than maximizing the biomass yield.

**Figure 3:**
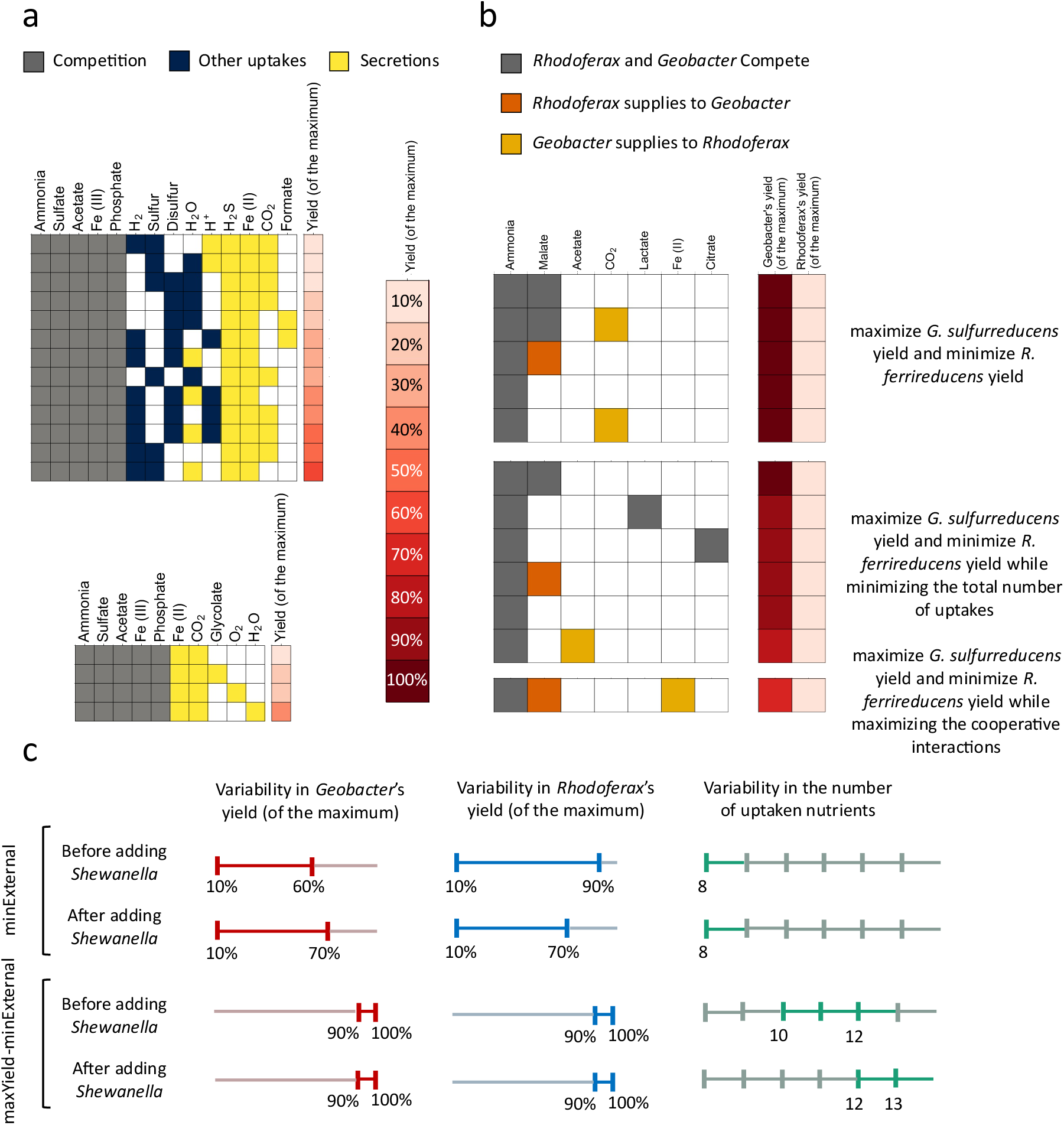
Various types of analyses on the uranium-reducing community. **a** Alternative DiMEs for *G. sulfurreducens* and *R. ferrireducens* that give rise to the most consistent interactions with the observations. In addition to the observed competition over ammonium, Fe (III), and acetate, the optimal solution includes obligatory competitions over sulfate and phosphate. Thirteen alternatives for *G. sulfurreducens* with yields ranging between 10% and 60% of the maximum yield and four alternatives for *R. ferrireducens* with yields ranging between 10% and 40% of the maximum yield could lead to the optimal interaction, **b** The optimal interaction patterns observed using Engineering objective functions, where the objective function was to maximize *G. sulfurreducens* yield and minimize *R. ferrireducens* yield, to maximize *G. sulfurreducens* yield and minimize *R. ferrireducens yield* while minimizing the total number of uptakes, and to maximize *G. sulfurreducens* yield and minimize *R. ferrireducens* yield while maximizing the cross-feeding interactions. Competition over ammonium, phosphate, and sulfate appeared in all cases, while the repeated presence of malate in the interactions indicates its importance in driving the community toward the desired state, and **c** Comparing the yields and the number of external uptakes before and after adding *S. oneidensis*, while minimizing the number of external uptakes, and maximizing the yield of all species while minimizing the total number of uptakes.

### Proposing synthetic media to achieve desirable properties

The third study was aimed at designing a community with desirable properties. Considering that only *G. sulfurreducens* remediates uranium contamination, we generated interaction networks in which *G. sulfurreducens* outperformed *R. ferrireducens* (i.e., *G. sulfurreducens* achieves a higher biomass yield than *R. ferrireducens*). To this end, we used three objective functions (Table 1). The first objective function selected the DiMEs with higher yields for *G. sulfurreducens* and the DiMEs with lower yields for *R. ferrireducens* (see Methods). The optimal solution was obtained when *G. sulfurreducens* grew with the maximum yield, and *R. ferrireducens* grew with the lowest possible yield, i.e., 10% of the maximum yield. We found one optimal interaction network (Figure 3b(i)).

In addition to favoring higher yields for *G. sulfurreducens* and lower yields for *R. ferrireducens*, the second and third objective functions minimized the number of uptaken nutrients and maximized the cross-feeding interactions, respectively (see Methods). The former resulted in five optimal interaction networks, while the latter had six optimal solutions (Figure 3b). Three interactions were essential using all the objective functions: competition over ammonium, sulfate, and phosphate. Interestingly, interaction through malate was observed in various alternatives more than other carbon sources, stressing the importance of this carbon source to induce the favorable phenotype in the community. Experiments can be performed to investigate the impact of providing malate to the community as a potential method to improve the uranium remediation capacity of the community.

### Adding *Shewanella oneidensis* enhances the uranium-reducing capacity of the community

In the final simulation, we used ReMIND to capture the impact of adding a new member on the interaction networks^42^. To this end, we chose *Shewanella oneidensis*, a bacterium with uranium-reducing capacity. Like *G. sulfurreducens*, *S. oneidensis* can reduce U (VI) to U (IV) and contribute to bioremediation. The impact of adding *S. oneidensis* to the community of *G. sulfurreducens* and *R. ferrireducens* was computationally investigated elsewhere^41^, where it was presumed that *S. oneidensis* consumes lactate and secretes acetate, which *G. sulfurreducens* and *R. ferrireducens* can then consume. Also, it was presumed that *S. oneidensis* competes with the other two species for Fe (III) and ammonium. These assumptions are based on observations about *S. oneidensis* in monoculture or other environments^43,44^. We used ReMIND to systematically examine the effect of this perturbation on the interaction network. We simulated the community with and without *S. oneidensis*, optimizing the Evolutionary objectives (Table 1) that resulted in the closest patterns to the observed interactions in the community of *G. sulfurreducens* and *R. ferrireducens* (Figure 2) (i.e., minExternal and maxYield-minExternal).

When the objective function was minExternal, the number of external resources remained unchanged after adding *S. oneidensis* (Figure 3c). The species either increase the overlap in their nutritional requirements through competitive interactions or increase their dependence on the secreted metabolites through cross-feeding interactions to reduce the number of external resources. In particular, in all optimal solutions for the two-member community, we observed either competition over the carbon source or cross-feeding. In both scenarios, a single external carbon source was used by the community. The interaction over the carbon source was mediated by four metabolites, i.e., malate (both competition and cross-feeding), citrate (only competition), lactate (only competition), and acetate (only cross-feeding).

Similarly, only a single carbon source was obtained from the environment, and two interactions via carbon sources were observed in all optimal solutions for the three-member community (competition or cross-feeding). The interactions were mediated by six metabolites, i.e., malate (both competition and cross-feeding), fumarate (both competition and cross-feeding), citrate (competition only), lactate (competition only), pyruvate (cross-feeding only), and succinate (cross-feeding only). The highest yields that *G. sulfurreducens* and *R. ferrireducens* could achieve were also affected; while the highest yield for *G. sulfurreducens* increased, the highest yield for *R. ferrireducens* decreased. This suggests that if the efficient consumption of resources is the sole factor in determining the interactions, the presence of *S. oneidensis* can benefit *G. sulfurreducens* and harm *R. ferrireducens*.

When we considered the second objective function, i.e., maxYield-minExternal, the yields of *G. sulfurreducens* and *R. ferrireducens* were unaffected, and both organisms could still achieve the highest possible yields (Table S1). However, the number of external resources increased after adding *S. oneidensis*. This implies that if diverse resources are available to the community, *G. sulfurreducens* and *R. ferrireducens* yields remain unaffected by *S. oneidensis*. Since we concluded in the previous sections that this community evolved under constrained nutrient availability in its natural environment, we could assume that adding *S. oneidensis* improves the bioremediation capacity both through its own uranium-reducing capability and by improving the yield of *G. sulfurreducens*.

### Closely related species have high nutritional overlaps

The gut microbiota of the honeybees shows numerous similarities with the human gut microbiome; thus, it has been widely used as a model system in microbiome research^45^. The core microbial community comprises five bacterial clusters, which are present in every female worker bee across the world^39^. These species include *Snodgrasella alvi* and *Giliamella apicola* from the Proteobacteria phylum, *Bifidobacterium asteroides,* which belongs to the Actinobacteria phylum, and lastly, *Lactobacillus* Firm-4 and *Lactobacillus* Firm-5 clades belonging the Firmicutes phylum. We chose *Lactobacillus mellifer*, *Lactobacillus mellis*, *Lactobacillus kullabergensis*, and *Lactobacillus apis* as representative species for the *Lactobacillus* Firm-4 and *Lactobacillus* Firm-5 clades, respectively^46^. Overall, we considered a seven-member core honeybee gut community for our analysis. In this study, we used the metabolic models from the CarveMe database to demonstrate our workflow’s adaptability with automatically reconstructed models^27^.

We defined a broad extracellular environment comprising 88 carbon-containing compounds (inorganic compounds were discounted) to generate DiMEs. This environment included the metabolites that could be utilized as substrates or secreted as by-products for all species, and they could originate from the host or microbiome metabolism (See Methods). We generated the DiMEs for all seven core members of the honeybee gut microbiome for ten different biomass yield regimes (Table S2). The alternatives varied among the species, with *Giliamella apicola* having the highest number of unique DiMEs across all seven species. In total, 86 metabolites were covered by these alternatives across all species, either as substrates or by-products. Analysis of these metabolites indicated that 36 could serve as substrates and by-products, 46 were only substrates, and 4 were only products (Figure 4a).

**Figure 4:**
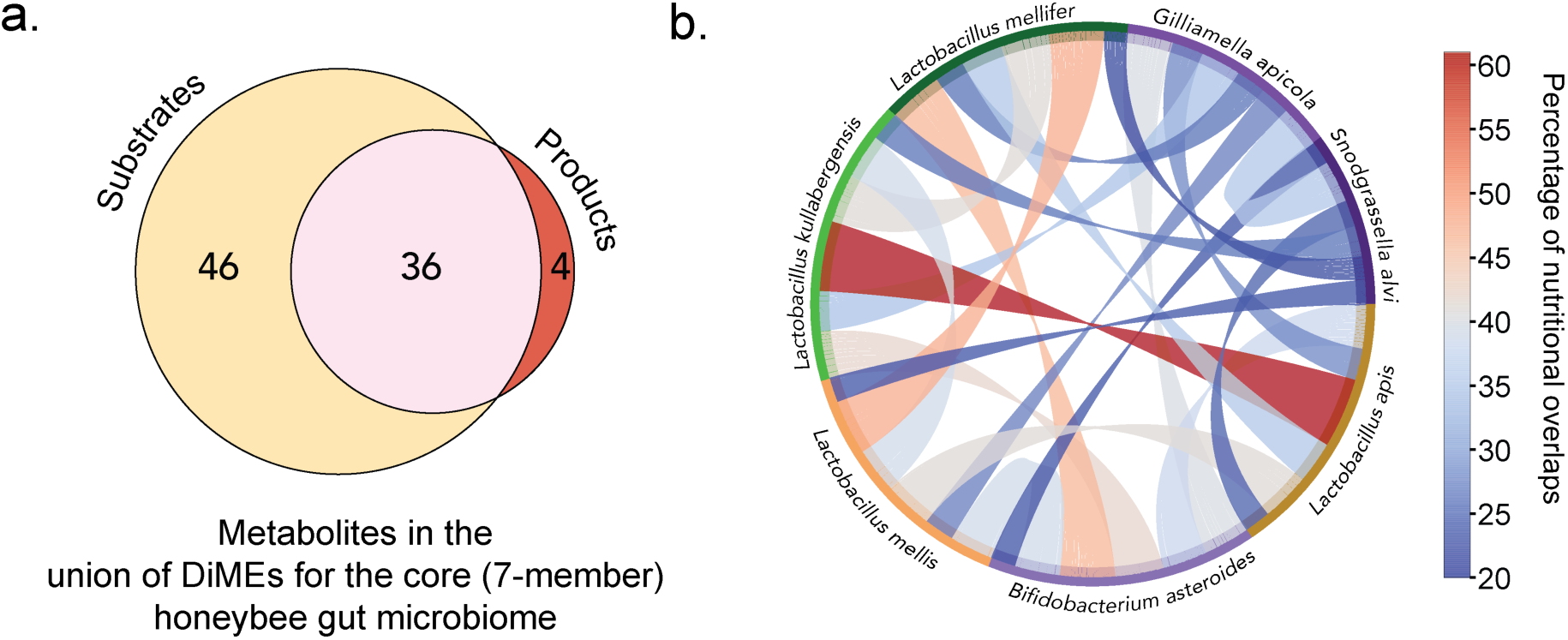
DiME analysis for the core honeybee gut microbiome. **a** The total number of metabolites that can be utilized as a substrate or secreted as a product from the union of DiMEs for all species and all biomass yield regimes. **b** Percentage of the nutritional overlaps between 21 pairs in the core honeybee gut microbiome.

We then identified the potential nutritional overlaps between every pair of species in the honeybee gut microbiome (Figure 4b) without differentiating between the alternatives. Therefore, the percentage does not refer to a particular physiology or habitat (See Methods). It instead reflects the similarity between the substrate utilization capabilities of both organisms and provides a comprehensive list of the potential competitive interactions. The highest nutritional overlap was between *L. kullabergensis* and *L. apis* from the *Lactobacillus* Firm-5 clade, with 34 overlapping nutrients across their DiMEs, leading to a 61% nutritional overlap. The next notable overlap was 48% between *L. mellifer* and *L. mellis* from the *Lactobacillus* Firm-4 clade. The high percentage predicted between the two Firmicutes clades supports the common belief that the closely related species are expected to have high nutritional overlaps and are primarily characterized by competitive interactions^26,27^. In contrast, the lowest percentage was 20% and found between *S. alvi* and *L. mellifer*. Comparing the nutritional overlaps for all the other species showed that *S. alvi* was dissimilar to the other community members in terms of substrate utilization, with the highest predicted nutritional overlap of 35% with *G. apicola* (Figure 4b). This dissimilarity was not surprising as most members of the honeybee gut are primary fermenters except for *S. alvi*, which has a significantly different metabolism^47,48^.

### Cross-feeding interactions become costly as the number of exchanged metabolites increases

We used ReMIND to identify the interaction network between *S. alvi* and *G. apicola*. These species, co-located in the ileum’s epithelial surface, exhibit complementary metabolisms: *G. apicola* ferments sugars, and *S. alvi* oxidizes carboxylic acids^47^. There has been increasing evidence for the cross-feeding interactions between the two species^47–49^. Therefore, we used the ReMIND framework to enumerate alternative directional cross-feeding patterns ranging from a single cross-fed metabolite to the maximum number of cross-fed metabolites in different environmental and biomass yield conditions.

Our study revealed 16 potential cross-fed compounds between the two species, including amino acids, carboxylic acids, and purine derivatives. We found that nine metabolites could be shared bidirectionally in the case of single-metabolite cross-feeding. In contrast, dihydroxy acetone, succinate, formate, acetate, and L-threonine, were provided unidirectionally by *G. apicola* to *S. alvi*, while 4-aminobutanoate and L-proline were provided by *S. alvi* to *G. apicola* (Figure 5a). These findings align with previous observations of specific cross-feeding interactions from *G. apicola* to *S. alvi* through lactate, formate^47^, succinate^49^, and 2-oxoglutarate^48^. The range of biomass yields for these interactions was wide, ranging from 10% to the optimum (100%) for both species, reflecting diverse metabolic costs and suggesting that some metabolites can be costlessly secreted^50^.

**Figure 5:**
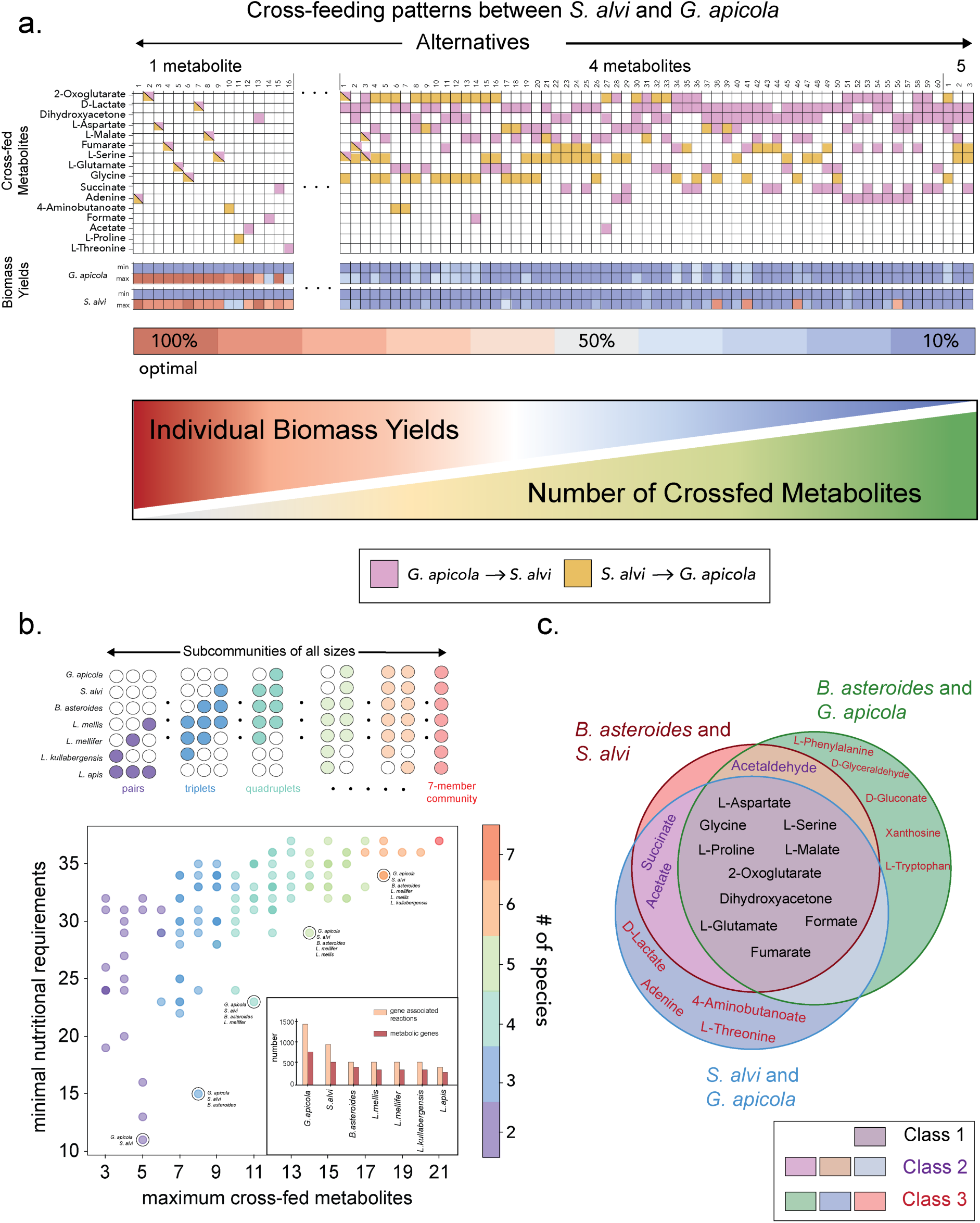
Subcommunity analysis in the core honeybee gut microbiome. **a** Analysis of the cross-feeding interactions between *S. alvi* and *G. apicola*. All alternative directional cross-feeding patterns between two species, from single compound (left) to maximal number of compounds exchanges (right), along with the minimal and maximal biomass yields predicted for each alternative interaction pattern. For visual purposes, the cross-feeding patterns for single, four and five metabolites exchanged are shown, dots represent the axis break, and the full analysis including the patterns with two and three metabolites exchanged can be found in Figure S1. **b** Characterization of all subcommunities of all sizes in the core honeybee gut microbiome according to the maximum cross-fed metabolites and minimal nutritional requirements. Circled communities indicate the sub-communities with the minimal nutritional requirements from each size; in general, their addition follows the order of species with decreasing number of metabolic genes. (Lower right corner). **c** Venn diagram of decomposed cross-feeding interactions (in the 3-member community) to their pairwise contributions, with their corresponding classes explained according to the number of pairs the metabolite can be cross-fed.

We predicted 90 cross-feeding patterns involving two metabolites. However, not all combinations of the 16 metabolites were feasible due to physiological and biochemical limits (Figure S1). These patterns fell into three categories: (i) *G. apicola* provided nutrients to *S. alvi* (23 cases), (ii) *S. alvi* provided nutrients to *G. apicola* (3 cases), and (iii) *G. apicola* and *S. alvi* mutually provided nutrients (64 cases). The first two classes represent commensalism or parasitism, while the latter represents mutualistic interactions. Biomass yield in these interactions varied from 10% to the optimum for both species, with mutualistic patterns typically leading to higher yields (Figure S1).

In the case of three-metabolite cross-feeding, we found 162 patterns. Of these, 134 represented mutualism, while 28 could be associated with commensalism or parasitism. Biomass yields for *G. apicola* varied from 10% to the optimum, and for *S. alvi*, from 10% to 90% of the optimum. Interestingly, the maximum yield for the nutrient provider in commensal or parasitic interactions is much lower, down to 10% for *S. alvi* and 40% for *G. apicola* (Figure S1). When four metabolites were exchanged, 60 patterns emerged, with 50 indicating mutualism and 10 commensalism or parasitism. Here, the maximum yield for *G. apicola* falls to 40% of the optimum, while *S. alvi* maintained up to 90%. Notably, *S. alvi* achieved higher yields when providing fewer or no nutrients to *G. apicola*.

For the maximum cross-fed metabolites, i.e., five cross-fed metabolites, we identified three patterns (Figure 5a). All patterns were mutualistic, where both species provided nutrients. In particular, *S. alvi*’s yield dropped to 10% of its optimum, while *G. apicola*’s maximum yield decreased to 30%. The overall trend observed across all patterns from one to five exchanged metabolites suggested that as the number of cross-feeding interactions increased, the highest individual biomass yields that could support these interactions decreased, indicating that exchanging metabolites becomes more costly for the species. In other words, if a species provided more nutrients, it did so at the expense of its biomass yield, reflecting apparent altruistic behavior.

### The same minimal environment can give rise to a variety of microbial interactions

We then assessed the nutritional needs for the two-member community consisting of *S. alvi* and *G. apicola*. To this end, we minimized the number of nutritional requirements supporting the growth of both species. This way, the community was expected to rely less on the habitat/host and more on the metabolic interactions. Our analysis showed that at least 11 nutrients must be externally provided (abiotic sources or host-derived metabolites) to the community to support the growth of both organisms. We generated all alternative minimal nutritional environments and the minimum and maximum individual biomass yield regimes that can be achieved within these environments (Figure S2a). Our study identified nine alternative minimal environments, each composed of 11 metabolites. These alternatives included 16 metabolites, containing amino acids, purines, pyrimidines, carboxylic acids, carbohydrates, and vitamins. Interestingly, our analysis showed that the same minimal environment can give rise to alternative interaction patterns, suggesting that the same environmental conditions can support diverse ecological interactions (Figure S2b and Supplementary Results).

### Metabolic interaction potential increases with the number of species in the microbial community

To examine the effect of changing the size of the community on the number of metabolic interactions, we reconstructed all subcommunities of the core honeybee gut microbiome of all sizes (Figure 5b). Our results showed that increasing the number of species in the random subcommunities increased both the maximum number of metabolites that can be exchanged and the minimal nutritional requirements. In addition, the variation in minimal nutritional requirements within the subcommunities of the same size decreased as the number of species increased.

We observed that within the subcommunities of the same size, species with a higher number of metabolic genes generally had the lowest number of nutrient requirements (marked in Figure 5b), consistent with previous findings for individual species^51^. Overall, it could be concluded that such communities relied less on the environment compared to subcommunities of the same size. To provide a lower and upper bound on the extent of competitive interactions that could occur, we also quantified the minimal and maximal competitive interactions (counted with the number of metabolites for which there is competition) within all subcommunities (Supplementary File 1). In accordance with the pairwise nutritional overlap (Figure 4b), the highest lower-bound for competitive interactions was observed for the *L. apis* and *L. kullabergensis* pair with at least 15 competitive interactions. Similarly, this value reached 11 competitive interactions for the *L. mellis* and *L. mellifer* pair.

### Examining the effect of biotic perturbations and identifying metabolite-hubs with high-connectivity

We next assessed how adding a new species (*B. asteroides*) to a two-member community (*S. alvi* and *G. apicola*) affects its minimal nutritional needs and cross-feeding interactions, an experiment known as N+1 biotic perturbation^52^ (see communities marked in Figure 5b). Our findings demonstrated that the external nutrient requirements increased from 11 to 15. We further analyzed the three-member community’s minimal environment by exploring different minimal environments and their impact on biomass yields (Figure S3 and Supplementary Results). Moreover, including *B. asteroides* increased the maximum number of cross-fed metabolites from five to eight (Figure 5b). In addition to the 16 metabolites that could be exchanged between *S. alvi* and *G. apicola*, six new metabolites contributed to cross-feeding interactions, including carboxylic acids, monosaccharides, organooxygen compounds, and amino acids. We then classified these metabolites based on the species exchanging them and decomposed the complex interaction networks into their pairwise contributions (Figure 5c).

Our classification of cross-fed metabolites yielded three classes. The first class contained ten metabolites, primarily amino acids and carboxylic acids, that could be exchanged between all species, thus linking the entire community. These metabolites mediate more robust interactions against biotic perturbations since interactions through these metabolites persist if a species is removed (Class 1 in Figure 5c). The second group included three metabolites that could only be exchanged due to the metabolic synergy of a particular species with the others. For instance, succinate could be only provided by *B. asteroides* and *G. apicola* to *S. alvi,* implying that such interaction emerges only if *S. alvi* is present (Class 2 in Figure 5c). Metabolites in both the first and second groups potentially serve as metabolite hubs, enhancing community connectivity beyond pairwise interactions. The last category of metabolites included nine compounds that could only be exchanged between a specific pair, representing only pairwise interactions without leading to higher degrees of connections (Class 3 in Figure 5c).

### Cross-fed metabolites in the core honeybee gut microbiome show different degrees of connectivity

We analyzed the seven-member core honeybee gut microbiome, focusing on the minimal nutritional environments and cross-feeding patterns, where the maximum number of metabolites was exchanged. This resulted in predicting 27 distinct minimal environments, each including 37 nutrients (Figure S4a). Our analysis showed that amino acids are the major components in these environments (Figure S4b, Supplementary Results). Regarding the cross-feeding interactions, we identified 93 patterns of 21 cross-fed metabolites (Figure 6a). The alternative cross-feeding patterns covered 29 metabolites, including mainly carbohydrates and carbohydrate conjugates, amino acids, and carboxylic acids (Figure 6b), consistent with previous research in diverse microbial ecosystems^27,50,53,54^. Ten of the 29 metabolites were identified as essential cross-fed metabolites, appearing in all alternatives of the maximum number of cross-fed metabolites.

**Figure 6:**
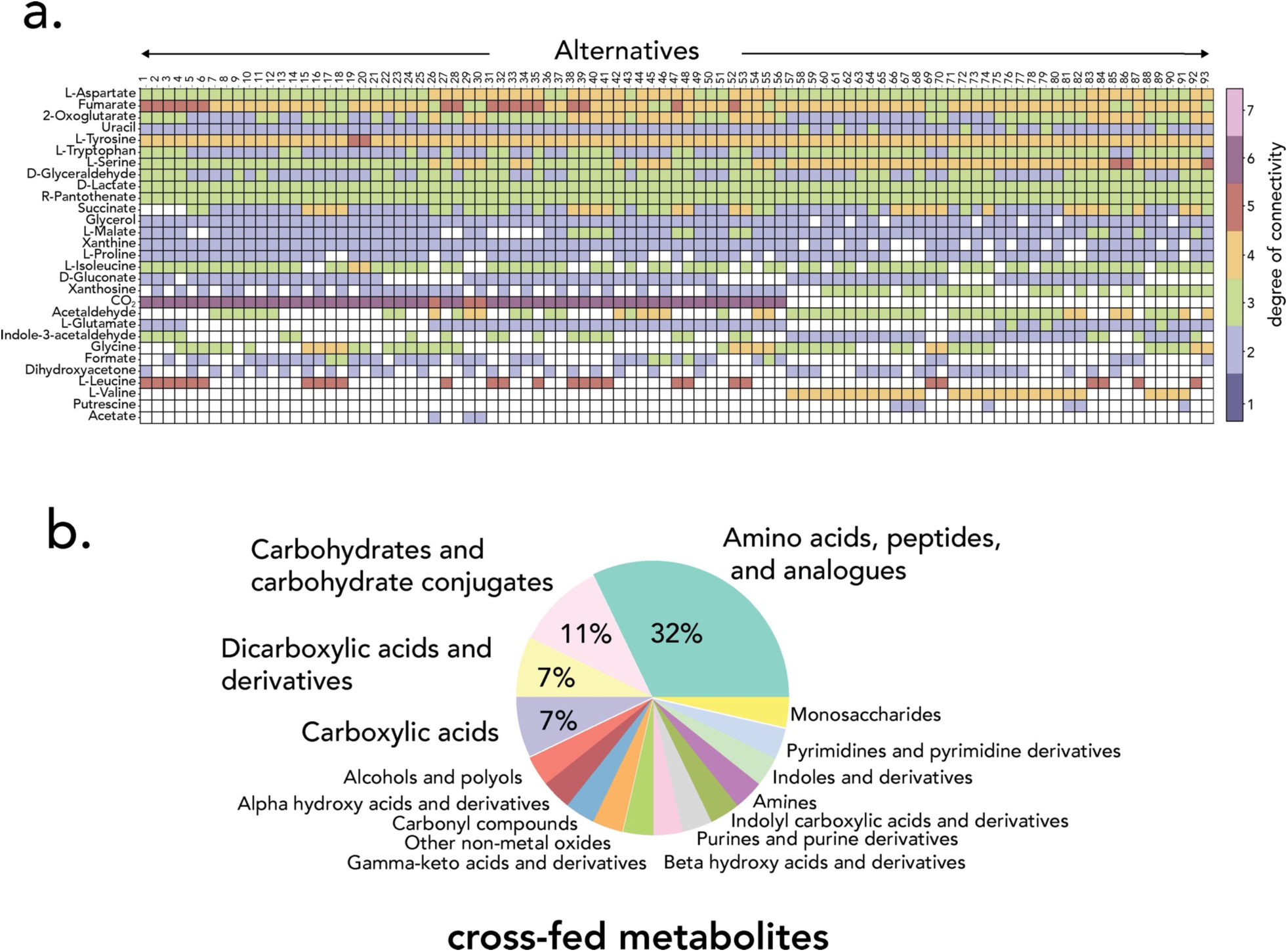
The analysis of the core honeybee gut microbiome. **a** Alternative cross-feeding patterns in the core microbiome when the maximal number (21) of metabolites are cross-fed, metabolites are colored according to the degree of connectivity of the metabolite (influx edges and outgoing edges). **b** Composition of the cross-fed metabolites in **a**., compounds are classified according to the Human Metabolome Database (HMDB), with 28/29 compounds classified (classification file taken from Machado et al.27).

We also assessed the degree of connectivity of metabolites in various patterns. To this end, we summed the maximum number of species that could uptake and produce the metabolite, representing the maximum number of links (edges) the metabolite could have. Our results indicated that different metabolites could provide different degrees of connectivity for the species. Carbon dioxide emerged as the cross-fed metabolite with the highest degree of connectivity, as all species can secrete CO_2_ in most, if not all, DiMEs, and only one species, *L. kullabergensis*, was predicted to take up small amounts of CO_2_ (not the primary carbon source) in some alternative DiMEs. A high degree of connectivity was also achieved by amino acids and carboxylic acids, including leucine, tyrosine, fumarate, and succinate. Interestingly, when the number of cross-fed metabolites was maximized, most of the exchanged metabolites had a connectivity degree of two. This implies that these compounds were only exchanged between a pair of species without leading to broader network connectivity (Figure 6a).

## Discussion

In this study, we presented our two-step framework for reconstructing microbial interaction networks using genome-scale metabolic models. In the first step, we formulated a MILP problem to generate the substrate and by-product profiles ranked by biomass yields for each species, which we named DiMEs. In the second step, we devised an ILP formulation to select a DiME for each species such that an objective function of interest is optimized. We benchmarked our method by generating interaction networks capturing experimentally observed interactions for a well-studied two-species uranium-reducing community^36,37^. We investigated ecological principles that shape interactions in this community, identified environments that support partial observations, and suggested potential interventions to enhance its bioremediation capacity, including the introduction of a new species to the community. We then applied our framework to the core honeybee gut microbiome composed of seven species. Consistent with previous studies^26,27^, we showed that closely related species have higher overlap in their nutritional requirements and are characterized mainly by competitive interactions. We demonstrated that our approach facilitates the analysis of metabolic interdependencies in different environments, providing insights into the trade-off between metabolite exchange and biomass yield. Additionally, it allows us to simplify complex interactions into simpler pairwise contributions, identify metabolite hubs with high connectivity, and assess the effect of adding or removing a member from a community.

Our framework offers several advantages for identifying metabolic exchanges within microbial communities. First, it requires a limited number of necessary inputs, namely the GEMs for the species and, if available, a list of potential extracellular metabolites (*e.g.*, exometabolomics). Second, our framework does not assume growth optimality for individual species. This is particularly important given the altruistic behavior of some microbial species^55,56^ and considering that metabolic systems do not always operate in a fully optimal state, allowing the system to be more robust to perturbations. Furthermore, our workflow systematically decomposes the problem into two sequential steps, significantly reducing computational complexity. We first address the problem at the species level, allowing parallel processing of DiMEs for each species. In the next step, the number of DiMEs to be included for each species can be controlled by adding them using a heuristic approach, making the framework easily scalable to larger communities. Finally, the DiMEs formulation itself is a valuable resource that provides insight into the metabolic capabilities of each organism. It provides the research community with a new approach to resolve the uncertainty associated with substrate and by-product profiles in metabolic models.

While we used automatically reconstructed metabolic models in our study, these models might need further refinement to accurately represent organism-specific phenotypes^30^.The scope of our analysis is inherently linked to the accuracy of these models, and future studies can further curate the models in combination with experimental validation to improve predictive accuracy. In addition, as more data becomes available, the analysis presented here can be extended by integrating additional data, such as host diet and partially observed interactions, to further refine and constrain the alternative interaction networks.

Our framework systematically identifies, ranks, and analyzes potential metabolic interactions within microbial communities. It holds great promise for experimental design and microbial therapy by guiding which species or metabolites to target to restore dysbiosis and prevent or treat diseases. Beyond microbial interactions, the methods developed here can be adapted to study various cellular interaction networks, including host-microbe and tumor microenvironment interactions, offering broader applicability in biological research.

## Methods

### The genome-scale metabolic models

Manually curated GEMs for *Geobacter sulfurreducens*^57^ and *Rhodoferax ferrireducens*^36^ were obtained from the literature. It was proposed that the nitrogen sources for this community are inorganic compounds such as ammonium or atmospheric nitrogen^37^. Thus, we blocked the uptake of organic compounds that contain nitrogen, including amino acids. We also assumed that each organism could use a single carbon source at a time due to the catabolite repression. This assumption was formulated as an additional constraint while generating DiMEs. Moreover, the uptake of oxygen was blocked to simulate the anaerobic environment^37^. We used manually curated GEM for *Shewanella oneidensis*^58^ to investigate the impact of adding a member to the community.

We assumed anaerobic conditions for the honeybee gut lumen. *S. alvi* is an obligate aerobe that colonizes the epithelium of the ileum, and thus has access to the oxygen diffused by the epithelial cells^39^. The other organisms of the honeybee gut microbiome used in this study grow in the absence of oxygen. We simulated growth under anaerobic conditions, and for the cases where the models failed to simulate growth under anaerobic conditions, we gap-filled them (Table S3) using the NICEgame workflow^59^ and the universal CarveMe model^30^ as a reaction pool. Also, the *S. alvi* GEM was gap-filled (Table S3) to enable growth on minimal media. We realized that import of some inorganic compounds (Ca^+2^, Zn^+2^ and Mn^2+^) to the cytosol was coupled with the transport of carboxylic acids (e.g., citrate) in some of the models, which unrealistically made the carbon source essential for the model. To address this issue, we modeled the transport of these compounds via the abc system or proton symport (Table S3).

### Thermodynamic curation of the genome-scale models

The process of thermodynamically curating Genome-scale Metabolic Models (GEMs) involves integrating thermodynamic data, specifically the Gibbs free energy of formation and its estimation error, into the models. The following pipeline was used for this estimation: First, MetaNetX (http://www.metanetx.org) was used to annotate the GEM’s compounds with identifiers from various databases, including SEED^60^, KEGG^61^, CHEBI^62^, and HMDB^63^. Then, using Marvin (version 20.20, 2020, ChemAxon http://www.chemaxon.com), the structures of compounds (canonical SMILES) were converted into their major protonation states at pH seven, and MDL Molfiles were generated. These Molfiles, along with the Group Contribution Method, were employed to calculate the standard Gibbs free energy of formation and its estimation error for the compounds. The calculation of the thermodynamic properties of the metabolites allowed the addition of thermodynamic constraints for a considerable percentage of the reactions of each GEM (Table S4).

### Defining the extracellular environment and general considerations

We defined a set of potential carbon sources and available inorganics for the uranium-reducing community (Supplementary File 2). For the core honeybee gut community, we first defined a broad extracellular environment. Instead of using the union of all exchanges from all models, we described a shorter but inclusive list of metabolites that could be utilized as substrates or secreted as by-products by all species. Inorganic compounds were assumed to be always present in the environment, and thus, they were not included in the metabolite list.

We divided the carbon sources into primary and supplementary (Supplementary File 3). The primary carbon sources were metabolites that could serve as the main carbon source for any of the species based on previous studies^48,49^ and genomic evidence^39,47^. The supplementary carbon sources included metabolites that were less likely to serve as main carbon source for any of the species (e.g., vitamins, purines, pyrimidines, certain amino acids) but were found to be essential (uptake or production) for growth of at least one of the models. We then differentiated between the two different types of carbon sources by constraining the maximum uptake rate of the supplementary carbon sources to 1 mmol gDW^-1^ h^-1^. The primary carbon source list was generated iteratively over all species starting from a user defined initial list of primary carbon sources. For the first iteration, we used the initial list and we also consider catabolite repression (i.e., only *C_u,b_* metabolites from the primary carbon source list can be uptaken at the same time) and growth was simulated for each models. If any of the models could not grow given these constraints, we added more metabolites to the primary carbon source list and/or relaxed the catabolite repression constraint (e.g., *C_u,b_*+1 metabolites from the primary carbon source list could be uptaken at the same time) and simulated growth again (Table S5). This process was repeated until all the models could grow. The uptakes for glucose and fructose, which were part of the primary carbon sources, were blocked for the species *S. alvi* to avoid any unrealistic alternatives, as this species is known to have lost all the pathways responsible for the breakdown of glycolytic sugars^39^. In addition, secretions of citrate, isocitrate glucose and fructose were blocked for all species.

For the uranium-reducing and the honeybee gut microbiomes, the maximum secretion values for all carbon sources and the maximum uptake rate for the primary carbon sources and inorganic compounds were constrained to 25 mmol gDW^-1^ h^-1^to avoid unrealistic fluxes. The maximum secretion rate for inorganics was also set to 50 mmol gDW^-1^ h^-1^. For this reason, some lower biomass yield regimes were found to be infeasible for *L. mellifer*, *L. mellis*, *L. apis* and *L. kullabergensis* and *B. asteroides*. DiMEs were exhaustively generated for *G. sulfurreducens*, *R. ferrireducens*, and *S. oneidensis* considering ten yield regimes. The DiMEs were exhaustively enumerated for all species in the honeybee gut and all feasible biomass yield regimes, including minimal, minimal+1, and minimal+2 sizes. 𝜇^’^was set to 0.2 h*^−^*^1^ for all species in this study and different biomass yield regimes were defined based on the uptaken moles of carbon.

### Decomposed in silico Minimal Exchanges (DiMEs)

Finding DiMEs is formulated as a MILP problem. It adds new variables and constraints to the FBA^64^ or TFA^65,66^ formulation to generate the minimal sets of active exchange reactions, including uptakes and secretions. In contrast to previously published formulations^67,68^, the nutrient and by-product profiles are decomposed based on the biomass yield to explore optimal and suboptimal solutions. In the first step, we find the optimal biomass yield for an experimentally observed growth rate (𝜇^’^) subject to the availability of the extracellular metabolites. The available extracellular metabolites can be defined through experimental studies (e.g., exometabolomics). If no information about the extracellular metabolites is available, we can generate DiMEs assuming all extracellular metabolites in the model are available. To find the optimal biomass yield for a fixed growth rate, we minimize the nutrient uptake:

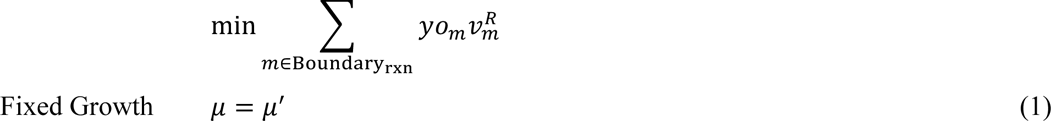

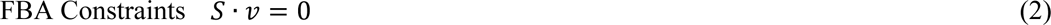

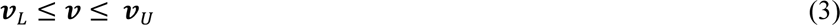

where 𝑆 is the stoichiometric matrix, 𝜇 the growth rate, 𝑣 are the reaction fluxes, and 𝑣*_L_* and 𝑣*_B_* are the lower and upper bound, respectively, for all the reactions in the network. In the objective function, 𝑦𝑜*_m_* represents a vector containing weights for the exchange reactions, where the exchange reaction *m* is linked to an extracellular metabolite denoted by *m*. The weighting vector is context-specific and can be defined by the user. We used the weighting vectors presented in Table 2. In addition to Equations (1)-(3), thermodynamic constraints can be considered to ensure thermodynamic feasibility^66,68^ (Equation (4)). 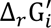 is the Gibb’s free energy of the reactions defined in TFA. 𝑏*^F^* and 𝑏*^R^* are the binary variables for the forward or reverse fluxes of all the reactions and M is a big constant.

**Table 2:** Different yield options to find and explore the optimal and suboptimal biomass yield space, where 𝑚 denotes each boundary reaction, and 𝑚, 𝑚𝑒𝑡 stands for the boundary metabolite associated with the reaction 𝑚. 𝑀𝑊, 𝑛_*C*_ and 𝑛_*N*_ denote the molecular weight of the 𝑚𝑒𝑡, the number of moles of carbon, and the number of moles of nitrogen in the metabolite 𝑚𝑒𝑡, respectively.

| Yield Option | Formula | Uses |
| --- | --- | --- |
| Gram uptake | $yo_m = MW_{m_{met}}$ | Penalizes large molecules,<br>Includes all extracellular metabolites (organic and inorganic). |
| Carbon uptake | $yo_m = n_{C_{m_{met}}}$ | When carbon is the limiting factor for growth, penalizes metabolites with a high carbon number |
| Nitrogen uptake | $yo_m = n_{N_{m_{met}}}$ | When nitrogen is the limiting factor for growth, penalizes metabolites with a high nitrogen number |

After finding the optimal biomass yield (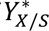), we fix the yield at various fractions of 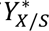 (referred to herein as yield regimes) by adding a new constraint. We also integrate new binary variables (𝑟*_m_*) to indicate if the exchange reaction *m* is active or inactive. The objective function is to maximize the number of inactive exchange reactions.

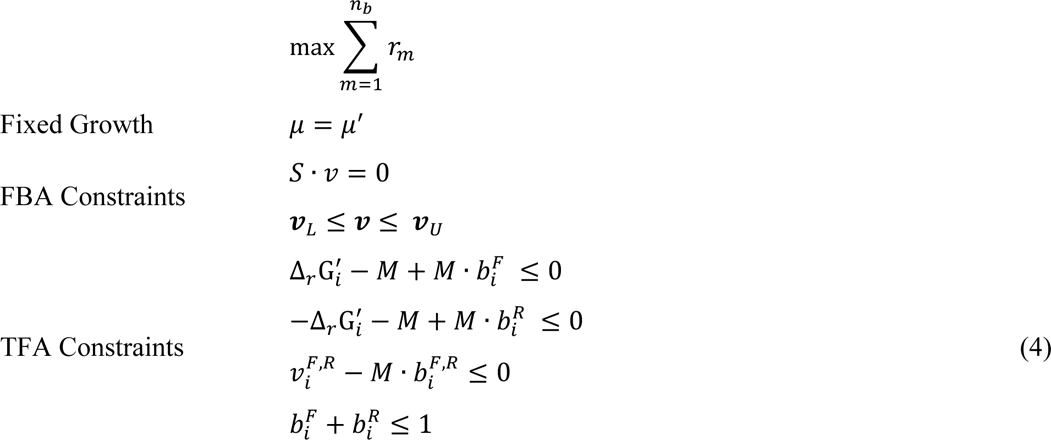

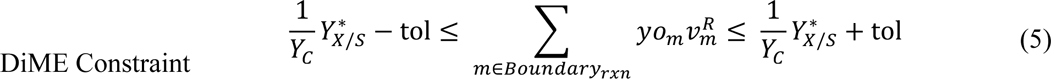

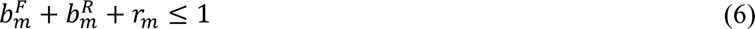

Equation (5) decomposes biomass yield into different yield regimes, where 𝑌*_C_* is a parameter defined as a fraction of the optimal biomass yield (e.g., 90%, 70%, 50%), and tol is a user-defined tolerance. The minimum and maximum yield regimes and the increments are user-defined parameters adjustable based on the user’s preference. If a DiME appeared in more than one yield regime, we only retained the highest yield regime, assuming that the organisms have evolved over generations to reach the highest possible yield in each environment. However, it is worth mentioning that this step is optional, and repeated DiMEs in different yield regimes can be retained. Equation (6) determines if an exchange reaction is active; if 𝑟_*m*_ = 1, the reaction *m* is inactive and does not carry flux, whereas if 𝑟_*m*_ = 0, the reaction *m* is active and carries flux.

We also included an additional constraint to account for catabolite repression:

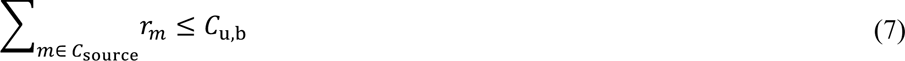

This constraint implies that the number of uptaken carbon sources must not exceed a user-defined parameter 𝐶_’,B_.

### Generating alternative DiMEs

To generate alternative nutrient and by-product sets, the following integer-cut constraint is added to the problem after generating each DiME (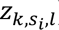) for the species 𝑠*_i_*.

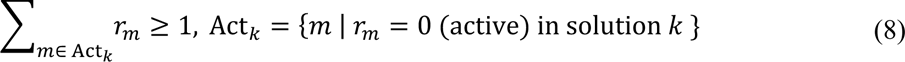

Since an exchange reaction *m* is only active if 𝑟*_m_* = 0 Equation (8) ensures that at least one of the exchange reactions is different in the next solution. We generate alternative solutions until a stopping criterion is reached. Stopping criteria can be set based on the number of alternatives, the size of DiMEs (e.g., minimal size, minimal size+1, minimal size+2), or exhaustive enumeration.

### Calculating nutritional overlap percentages

Nutritional overlaps between pairs were determined by calculating the percentage of shared substrates out of the total unique substrates for both members of each pair, considering all DiMEs across different biomass yield scenarios. This calculation is similar to the Jaccard index and is done as follows:

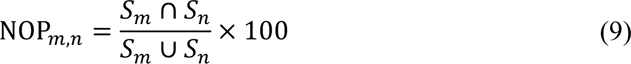

where *Sm* and *Sn* represent unique substrates of the species *m* and *n* respectively.

### Reconstruction of interaction network

The generated set of DiMEs includes all metabolic capabilities of a species given an extracellular environment, where each DiME represents a specific physiology. To represent the species with the most relevant DiMEs, we can optimize an objective function representing a community driving force, partially observed interactions, or an engineering goal. We devised an Integer Linear Programming (ILP) problem to select one DiME per species such that a user-defined objective is optimized. The ILP problem includes the following constraints:

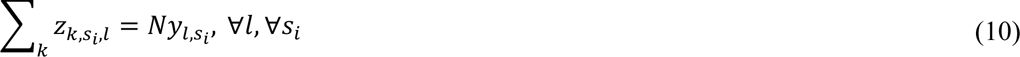

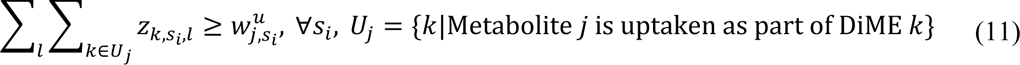

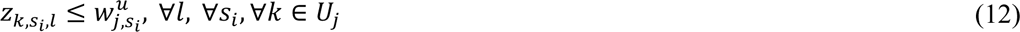

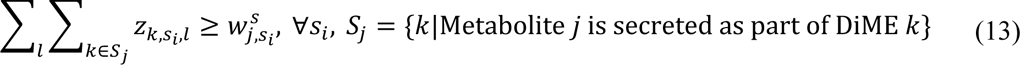

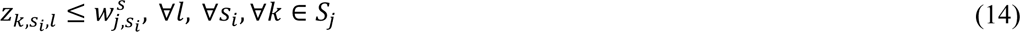

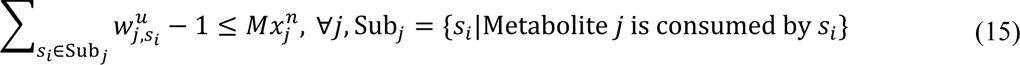

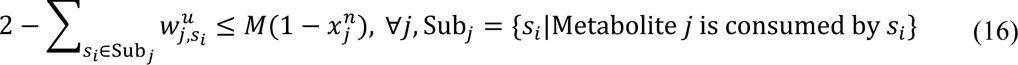

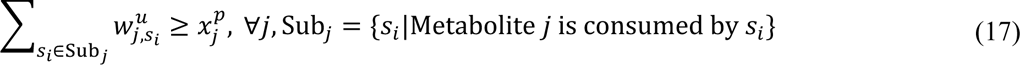

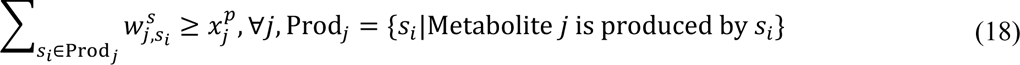

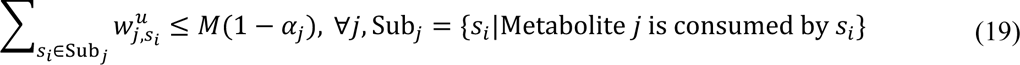

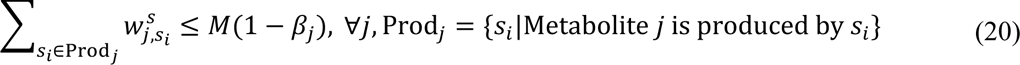

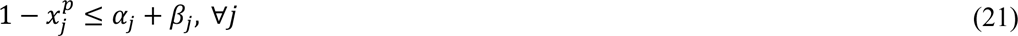

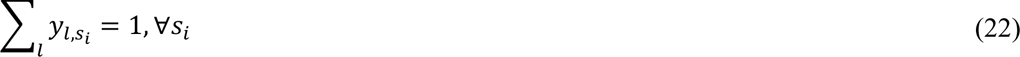

The variables, indices, and constants are described in Table *3*. Equation (10) defines the number of DiMEs selected per species for each yield regime. We used a default value of *N*=1 to select one DiME per species. However, to capture the heterogeneity in single species or to account for complex metabolisms, such as co-utilization of a nutrient or co-production of a by-product, higher values of *N* can be used to allow the selection of multiple DiMEs per species. Equations (11) and (12) enforce that the *j*th metabolite uptake is active if and only if at least one DiME is selected in which this metabolite is among the nutrients.

**Table 3:** The definition of variables, indices, and constants used in the formulation.

| Symbol | Type | Represents |
| --- | --- | --- |
| $i$ | Index | All reactions in the model |
| $m$ | Index | Boundary (exchange) reactions in the model |
| $m_{met}$ | Index | Boundary metabolite |
| $j$ | Index | Extracellular metabolite |
| $k$ | Index | DiME |
| $l$ | Index | Yield regime |
| $s_i$ | Index | Species |
| $r_m$ | Binary Variable | If the $j$th boundary reaction is active 0, else 1 |
| $x_j^n$ | Binary Variable | If competition over the $j$th metabolite exists 1, else 0 |
| $x_j^p$ | Binary Variable | If cooperation over the $j$th metabolite exists 1, else 0 |
| $w_{j,s_i}^u$ | Binary Variable | If the $j$th metabolite by the $s_i$th species is uptaken 1, else 0 |
| $w_{j,s_i}^s$ | Binary Variable | If the $j$th metabolite by the $s_i$th species is secreted 1, else 0 |
| $z_{k,s_i,l}$ | Binary Variable | If the $k$th DiME in the $s_i$th species and the $l$th yield regime is active 1, else 0 |
| $y_{l,s_i}$ | Binary Variable | If the $l$th yield regime in the $s_i$th species is active 1, else 0 |
| $\alpha_j$ | Binary Variable | If at least one species uptakes the $j$th metabolite 0, else 1 |
| $\beta_j$ | Binary Variable | If at least one species secretes the $j$th metabolite 0, else 1 |
| $N$ | Constant | The number of DiMEs that can be chosen per species |
| $M$ | Constant | An arbitrarily large number bigger than the number of species for the ILP and bigger than all flux upper bounds for TFA |

Similarly, Equations (13) and (14) ensure that the *j*th metabolite secretion is active if and only if at least one DiME is selected, in which this metabolite is among the by-products. Equations (15) and (16) account for the competition over the *j*th metabolite if at least two species in the community consume this metabolite. On the other hand, Equations (17)-(21) capture cooperation over the *j*th metabolite if at least one species produces this metabolite and one other species consumes it. Finally, Equation (22) ensures that only one yield regime is active for each species.

The above constraints are the complete ILP formulation to capture competitive and cross-feeding interactions independent of the objective function. However, if the objective function is to maximize or minimize the interactions, some of the abovementioned constraints can be removed to simplify the problem.

### Code and Data Availability

ReMIND is implemented in both Python and MATLAB. The Python version is based on pyTFA^65^, a Python implementation of the TFA method. It uses COBRApy^69^ and Optlang^70^. The MATLAB version is based on matTFA^65^, a MATLAB implementation of the TFA method. The code used to generate the models is freely available under the APACHE 2.0 license at https://github.com/EPFL-LCSB/remind.

Computations for the uranium-reducing community were done on 32-bit macOS, Intel® Xeon® CPU 2.7 GHz, 32 GB 1866 MHz RAM and for the honeybee gut community on 64-bit Ubuntu 20.04.2 LTS, Intel® Xeon® Gold 6254 CPU 3.10 GHz (18 cores, 36 threads per socket), 192 GB 3200 MHz RAM. Code was run on Python 3.6 on Docker (20.10.21) containers based on the official python 3.6-stretch container using the optlang package^70^ and using commercial solver ILOG CPLEX version 12.8.0.0.

## Acknowledgments

The authors would like to thank *Dr. Andrew Quinn* and *Prof. Philipp Engel* for the for the valuable discussions on the bee gut microbial community. Funding for this work was provided by the Swiss National Science Foundation (SNSF): grant 200021_188623, NCCR Microbiomes, a National Centre of Competence in Research (grant number 180575), the European Union’s Horizon 2020 research and innovation programme under grant agreement No 814408, and the École Polytechnique Fédérale de Lausanne.

## Author Contribution

OO, AS, and VH designed the study. OO and AS reviewed the literature and implemented the code in python. OO ran the simulations and performed thermodynamic curation, initial analysis, and visualization for the uranium-reducing community. AS ran the simulations and performed thermodynamic curation, initial analysis, and visualization for the bee community. EV performed the gap-filling for the uranium-reducing and the bee communities and implemented the code in MATLAB. OO, AS, and EV conducted the final analysis of the results. OO, AS, EV, and VH provided the discussion and wrote the manuscript. VH was responsible for acquiring resources, funding, project administration, and supervision.

