## Supplementary Information for "A constraint-based framework to reconstruct interaction networks in microbial communities"

**Supplementary Methods**

### The objective function to reconstruct the interaction networks

It is possible to use different objective functions with the ILP formulation. Here, we provide the mathematical formulation of the objective functions used in this study. The notations are the same as in Table 3 in the main text. The following expression was used to minimize the total number of uptakes:

|  | $\min_{} \sum_{s_{i}} \sum_{j} w_{j,s_{i}}^{u}$ |
| --- | --- |

An additional constraint was formulated to differentiate between the external metabolites and the metabolites that are produced and provided by other species:

$$w_{j,s_{i}}^{u}-x_{j}^{p}\leq u_{j}, \forall j, \forall s_{i}$$

(S1)

Then, the objective function is defined as follows:

|  | $\min_{} \sum_{j} u_{j}$ |
| --- | --- |

Since we minimize the sum of binary variables $u_{j}$ in the objective function, Equation (S1) enforces $u_{j}$ to be equal to 1 only if $w_{j,s_{i}}^{u}=1$ and $x_{j}^{p}=0$, which implies that the metabolite *j* is an external resource. We used $\min_{} \sum_{j} x_{j}^{n}$ to minimize the competitive interactions and
$\max_{} \sum_{j} x_{j}^{p}$ to maximize cooperative interactions. To account for the active yield regime in the species $s_{i}$, we used the following expression:

|  | $\sum_{l} {ly}_{l,s_{i}}$ |
| --- | --- |

Since only one yield regime is active (Equation 22), maximizing (minimizing) this expression results in selecting the highest (lowest) yield regime. Finally, the following objective function rewards activation of the observed interactions and inactivation of the interactions that are not observed to find the most compatible interaction network with the observations (for the uranium-reducing community):

$$\min_{} \left[ \sum_{j\in\{{NH}_{4}^{+},{Fe}^{3+},Acetate\}} {(1-x}_{j}^{n})+\sum_{j\notin\{{NH}_{4}^{+},{Fe}^{3+},Acetate\}} x_{j}^{n}+\sum_{j\notin\{{NH}_{4}^{+},{Fe}^{3+},Acetate\}} x_{j}^{p} \right]$$

### Generating alternative interaction networks

Similar to the DiMEs alternatives, alternatives are generated at the community level, based on interactions, cooperations, competitions, or different minimal environments. This is ensured based on the integer cuts added after generating a solution. The formulation of the integer cut depends on the interested alternatives that we want to generate for the given objective function. For example, the following integer cut is suitable to generate alternative cross-feeding patterns:

$$\sum_{j\in\mathrm{In}\mathrm{Act}_{h}} x_{j}^{p}+\sum_{j\in\mathrm{Act}_{h}} (1-x_{j}^{p})\geq1,$$

$$\mathrm{Act}_{h}=\left\{ j | x_{j}^{p}=1 \mathrm{in} h\mathrm{th} \mathrm{solution} \right\}, \mathrm{InAct}_{h}=\left\{ j | x_{j}^{p}=0 \mathrm{in} h\mathrm{th} \mathrm{solution} \right\}$$

(2)

This way, we ensure that the new solution is different in terms of cross-fed metabolites from the previously found solutions.

To generate new cross-feeding patterns while differentiating between the consumers and producers in the same pattern, we used the following integer cut:

$$\sum_{j\in\mathrm{InAct}_{h}^{1}} x_{j}^{p}+\sum_{j\in\mathrm{Act}_{h}^{1}} \left( 1-x_{j}^{p} \right)+$$

$$\sum_{s_{i}} \sum_{j\in\mathrm{InAct}_{h}^{1}} w_{j,s_{i}}^{u}+\sum_{s_{i}} \sum_{j\in\mathrm{Act}_{h}^{2}} \left( 1-w_{j,s_{i}}^{u} \right)+$$

$$\sum_{s_{i}} \sum_{j\in\mathrm{InAct}_{h}^{3}} w_{j,s_{i}}^{s}+\sum_{s_{i}} \sum_{j\in\mathrm{Act}_{h}^{3}} \left( 1-w_{j,s_{i}}^{s} \right)\geq1,$$

$$\mathrm{Act}_{h}^{1}=\left\{ j | x_{j}^{p}=1 \mathrm{in} h\mathrm{th} \mathrm{solution} \right\}, \mathrm{InAct}_{h}^{1}=\left\{ j | x_{j}^{p}=0 \mathrm{in} h\mathrm{th} \mathrm{solution} \right\},$$

$$\mathrm{Act}_{h}^{2}=\left\{ j | w_{j,s_{i}}^{u}=1 \mathrm{in} h\mathrm{th} \mathrm{solution} \right\}, \mathrm{InAct}_{h}^{2}=\left\{ j | w_{j,s_{i}}^{u}=0 \mathrm{in} h\mathrm{th} \mathrm{solution} \right\},$$

$$\mathrm{Act}_{h}^{3}=\left\{ j | w_{j,s_{i}}^{s}=1 \mathrm{in} h\mathrm{th} \mathrm{solution} \right\}, \mathrm{InAct}_{h}^{3}=\left\{ j | w_{j,s_{i}}^{s}=0 \mathrm{in} h\mathrm{th} \mathrm{solution} \right\}$$

(3)

Likewise, the following integer cut ensures that each new solution represents a new set of minimal environments:

$$\sum_{j\in\mathrm{In}\mathrm{Act}_{h}} u_{j}+\sum_{j\in\mathrm{Act}_{h}} (1-u_{j})\geq1,$$

$$\mathrm{Act}_{h}=\left\{ j | u_{j}=1 \mathrm{in} h\mathrm{th} \mathrm{solution} \right\}, \mathrm{InAct}_{h}=\left\{ j | u_{j}=0 \mathrm{in} h\mathrm{th} \mathrm{solution} \right\},$$

(4)

Finally, by including the binary variables for yield regimes, $y_{l,s_{i}}$, we can differentiate solutions based on the yield regimes.

**Supplementary Results**

### Minimal nutritional environments for *S. alvi* and *G. apicola*

#### Each alternative minimal environment consists of two classes of metabolites. The first class is the essential nutrients present in all nine alternatives, while the second class is the substitutable metabolites that alternate within alternative minimal environments. The essential metabolites comprise nine compounds, mainly amino acids and vitamins. The remaining seven metabolites can be grouped into two classes of compounds alternating across the different solutions and can substitute each other. The first class consists of the purines adenine and inosine, and the second class consists of the carbohydrates and derivatives sucrose, D-glucose D-fructose, and 5-dehydro D-gluconate and the carboxylic acid formate. Any combination of metabolites from these two alternating groups results in an alternative minimal environment, except for formate, which was identified as part of the minimal environment only when combined with adenine (Figure S2a).

#### In all alternative minimal environments, at most three out of the 11 metabolites were uptaken by both species, resulting in competition between *S. alvi* and *G. apicola* and most of the re- maining nutrients were only utilized by *G. apicola*. The lower biomass yield regimes observed for both species hint at metabolic cross-feeding between the species in these environments. To illustrate this, we have focused on one alternative (Alternative 7, marked (*) in Figure S2a) from the minimal environment and showed that the same minimal environment can give rise to different interaction patterns. More specifically, two different metabolic interaction patterns can be achieved with the same nutritional environment, with varying individual biomass yields (Figure S2b). Our analysis showed that in both patterns, *S. alvi* provides L-glutamate to *G. apicola* and *G.apicola* provides 2-oxoglutarate in return to *S. alvi*. The distinguishing factor lies in one pattern where *S. alvi* additionally provides D-lactate to *G. apicola*. Considering biomass yields, *S. alvi* can sustain both interaction patterns with the same biomass yield. However, for *G. apicola*, the biomass yield supporting the resulting interaction network increases when there is no cross-feeding involving D-lactate. In the high biomass yield scenario, *G. apicola* produces 2-oxoglutarate, L-malate, acetaldehyde, and carbon dioxide, while in the lower biomass yield regime, it produces L-aspartate instead of L-malate, also in addition to the previously available metabolites it also uptakes D-lactate which is provided by *S. alvi*. This shift suggests that the production of L-aspartate is more costly than the production of L-malate in this minimal environment for *G. apicola*.

### Biotic perturbations can alter the minimal environment

#### In contrast to the two-species community, we predicted a unique minimal environment, composed of 15 minimal compounds. The composition of the environment included 10 nutrients from the 2-species community and an additional five compounds that were specifically required for *B. asteroides* growth and were only uptaken by *B. asteroides* in the minimal environment (Figure S3). Interestingly, none of the five alternating carbohydrates and derivatives in the 2-species community were part of the minimal environment when *B. asteroides* was introduced, suggesting cross-feeding interactions for these sources. Our analysis showed that this minimal environment could support three different cross-feeding interaction profiles. In all profiles, formate, which was one of the alternating carbon sources in the 2-species community (Figure S2a), is provided to *G. apicola* by *B. asteroides*. Similarly, aspartate, glutamate and 2-oxoglutarate are part of all three interaction patterns, and acetate appears to be cross-fed from *B. asteroides* to *S. alvi* in only one of the patterns. In all three interaction patterns, *G. apicola* and *B. asteroides* maintained the same biomass yield, which was 80% and 30% of their optimal biomass yield, respectively. However, *S. alvi* achieved three different biomass yields for three interaction profiles.

### Amino acids constitute a major part of the essential nutrients for the core honeybee gut microbiome

#### We generated all alternative minimal nutritional environments that support the growth of the core honeybee gut microbiome. We predicted 27 alternative minimal nutritional require- ment profiles, with 37 metabolites in each alternative, indicating that at least 37 compounds are required to be supplied externally to the honeybee gut microbiome. In total, 27 alternatives covered 46 compounds, with 30 essential and 16 alternating nutrients (Figure S4a). Most of the essential nutrients comprise amino acids, which make up 43% of the essential com- pounds (Figure S4b). A major part of the alternating nutrients is composed of amino acids, purines and derivatives, and carbohydrates and their conjugates (Figure S4b). Furthermore, we classified the nutrients in the 27 alternative environments according to the number of species that uptake them. Our analysis shows that apart from 13 compounds, which were only uptaken by a single species in all alternatives, the remaining 33 compounds could be uptaken by two or more species, resulting in competitive interactions in the core honey bee gut microbiome. Our results indicate that all species require L-arginine and at least three species take up most of the amino acids in the environment. The essential carbohydrate present in all minimal environments was fructose, the most abundant sugar in the diet of the bees [1], and we predict that it is used by four to five species in the honeybee gut microbiome (Figure S4a). Lastly, we assessed the possible number of cross-feeding interactions when the core microbiome grows in minimal environments. The results indicate that this value ranges between 11 to 16 in these minimal environments, which is below the predicted number for maximal cross-feeding in this community. Overall, this suggests that minimization of dietary requirements and maximization of cross-feeding interactions do not lead to the same network of interactions for the core honey bee gut microbiome.

**Supplementary Figures**

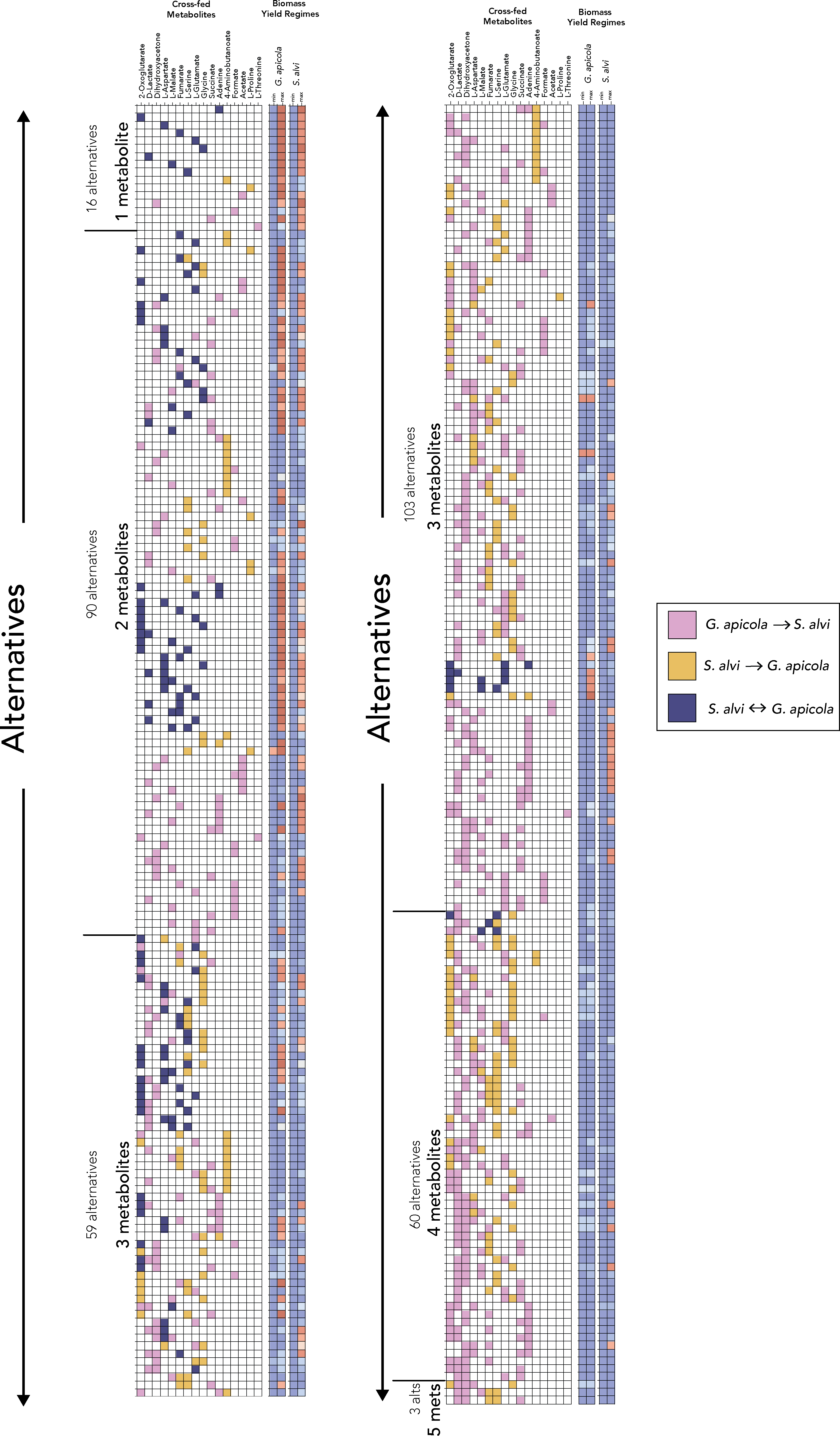

Figure S1: All alternative directional cross-feeding patterns between *S. alvi* and *G. apicola*. Colors represent the direction of the interaction; blue indicates that the alternatives were found where the metabolite can be exchanged both ways from *S. alvi* to *G. apicola* and from *G. apicola* to *S. alvi*.

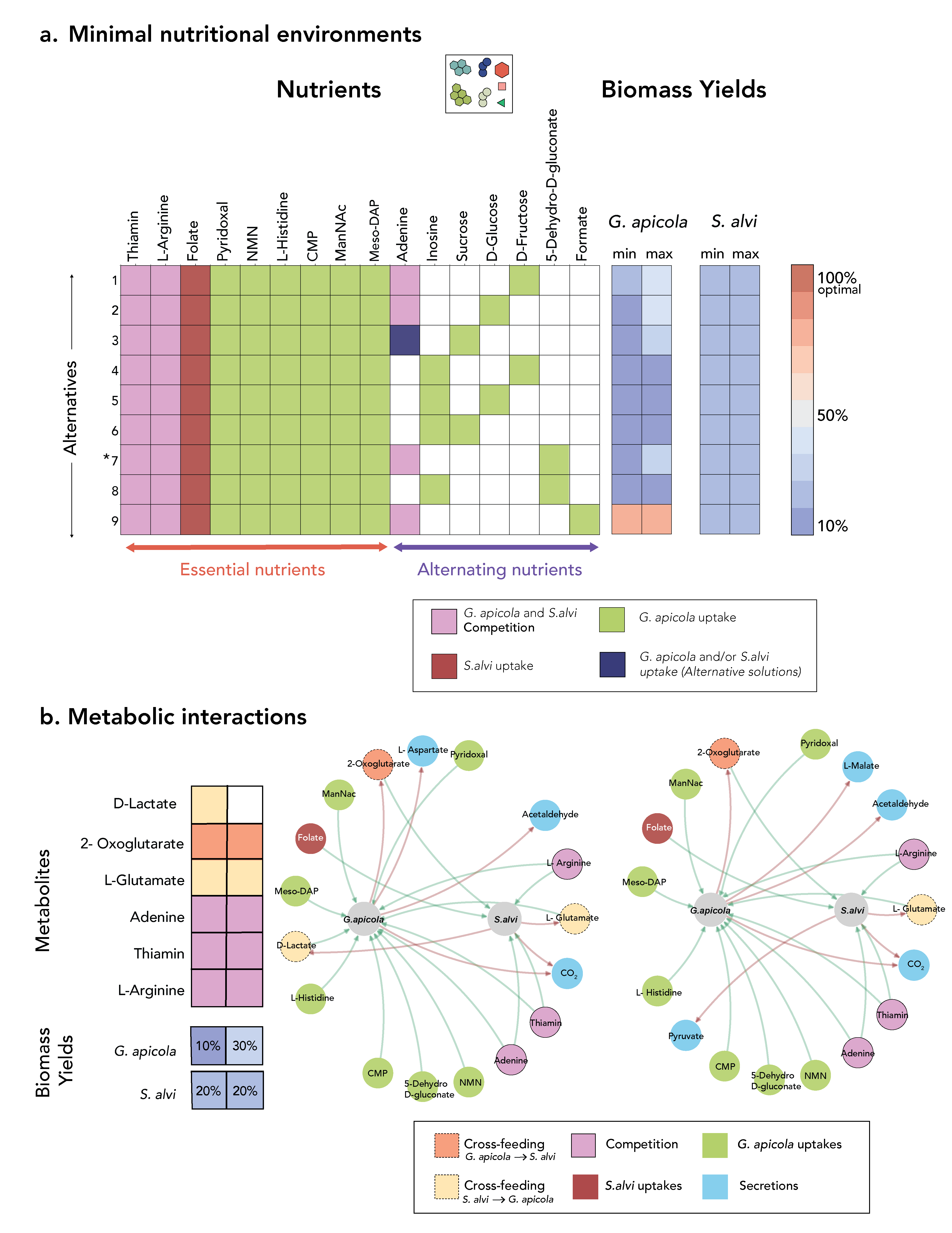

Figure S2:Minimal nutritional requirement analysis for the community of *S. alvi* and *G. apicola*. a) All alternative minimal nutritional environments that support the community’s growth for S. alvi and G. apicola with the min and max biomass yield regimes for each species. Nutrients are colored according to the species that uptake the nutrient, dark blue indicates that the nutrient uptake can vary at varying biomass yield regimes. b) Different interaction patterns and their resulting interaction networks visualized for the same minimal environment (Alternative 7 in a (*)), metabolites are colored according to the legend. In the legend, meso-2,6 diaminoheptanedioate, cytidine monophosphate, nicotinamide mononucleotide and N-acetyl-D-mannosamine are abbreviated as Meso-DAP CMP, NMN and ManNac, respectively.

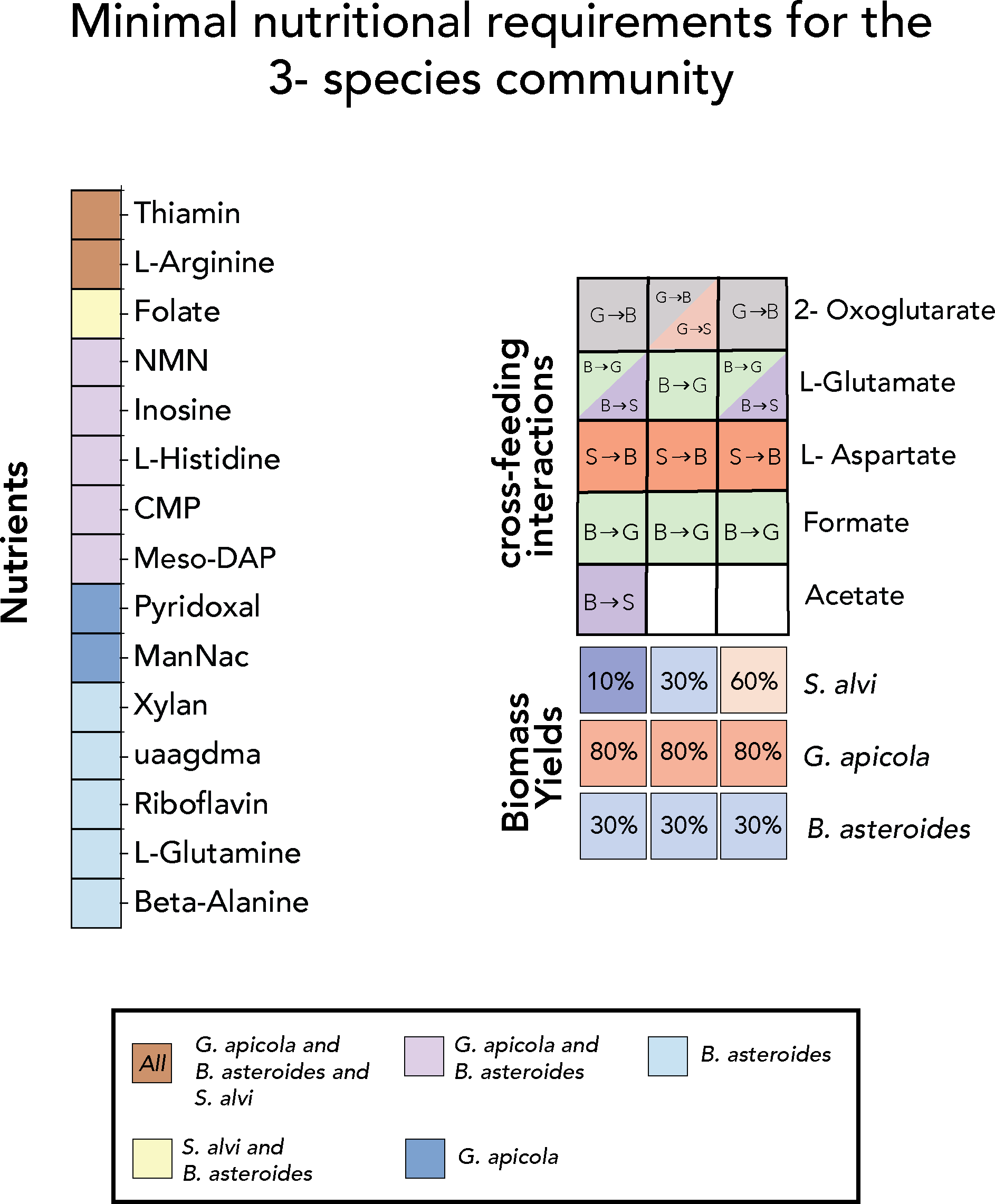

Figure S3: Minimal nutritional requirement analyses for the community *S. alvi*, *G. apicola*, and *B. asteroides*, nutrients are colored according to the species or combinations of species that uptake them (left). Different cross-feeding patterns observed in the minimal environment with the predicted biomass yields for each species (right). In the legend, meso-2,6 diamino- heptanedioate, cytidine monophosphate, nicotinamide mononucleotide and N-acetyl-D- mannosamine are abbreviated as Meso-DAP CMP, NMN and ManNac respectively.

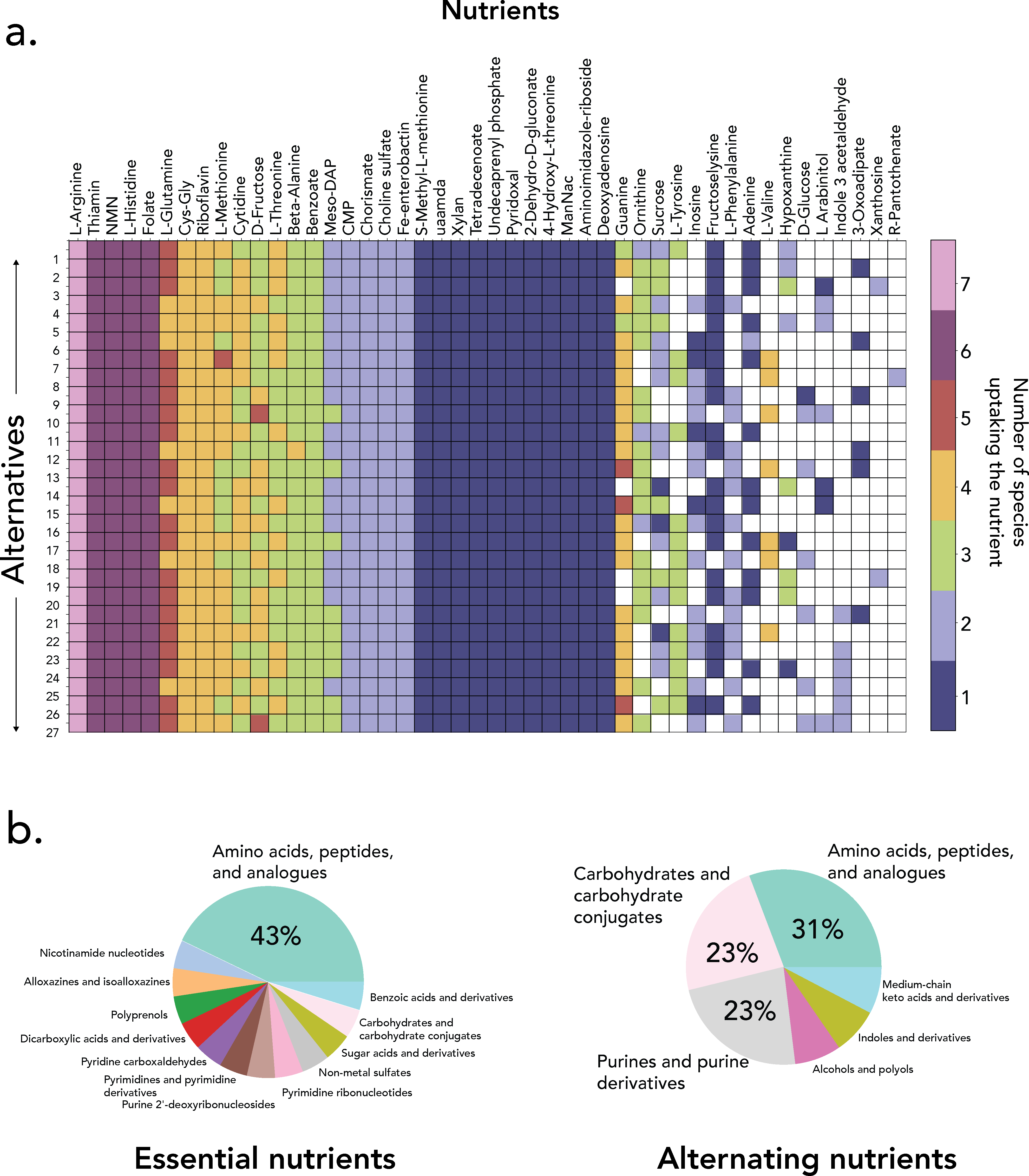

Figure S4: The analysis of the core honeybee gut microbiome with respect to the minimal nutritional requirements. a) All alternative minimal environments, each composed of 37 metabolites supporting the growth of the core honeybee gut microbiome. Metabolites are colored with respect to the number of species uptaking the nutrient in that environment. In the legend, eso-2,6 diaminoheptanedioate, cytidine monophosphate, nicotinamide mononucleotide and N-acetyl-D-mannosamine are abbreviated as Meso-DAP CMP, NMN and ManNac respectively. b) Composition of the minimal nutritional requirements grouped into essential nutrients and alternating nutrients, compounds are classified according to the Human Metabolome Database (HMDB), with 21/30 essential nutrients and 13/16 alternating nutrients classified (classification file taken from Machado et al. [2]).

**Supplementary Tables**

Table S1: The number of DiMEs for G. sulfurreducens, R. ferrireducens, and S. oneidensis at different yield regimes.

| Yield regimes (percent of the max. yield) | Number of DiMEs | | |
| --- | --- | --- | --- |
|  | *G. sulfurreducens* | *R. ferrireducens* | *S. oneidensis* |
| 10% | 162 | 215 | 273 |
| 20% | 172 | 175 | 336 |
| 30% | 186 | 151 | 170 |
| 40% | 173 | 109 | 96 |
| 50% | 146 | 94 | 32 |
| 60% | 97 | 57 | 7 |
| 70% | 70 | 28 | 2 |
| 80% | 40 | 17 | 1 |
| 90% | 18 | 10 | 1 |
| 100% | 1 | 1 | 1 |
| Unique DiMEs | 519 | 400 | 809 |
| Substrates | 18 | 19 | 16 |
| Products | 16 | 15 | 28 |

Table S2:The total number of generated DiMEs for the core honeybee gut microbiome and the statistics. Total number of metabolites, substrates, and products are calculated based on the union of all DiMEs for each species.

| **Biomass yield regimes** | ***G. apicola*** | ***S. alvi*** | ***L. apis*** | ***L.kullabergensis*** | ***L. mellifer*** | ***L.mellis*** | ***B.asteroides*** |
| --- | --- | --- | --- | --- | --- | --- | --- |
| 10% | 1346 | 2848 | - | - | - | - | - |
| 20% | 424 | 734 | - | 12 | - | - | 181 |
| 30% | 623 | 553 | 8 | 36 | - | - | 190 |
| 40% | 198 | 172 | 10 | 49 | - | - | 55 |
| 50% | 111 | 132 | 11 | 42 | - | - | 39 |
| 60% | 53 | 112 | 192 | 251 | 55 | 21 | 61 |
| 70% | 90 | 72 | 85 | 202 | 33 | 20 | 108 |
| 80% | 764 | 130 | 119 | 202 | 33 | 20 | 15 |
| 90% | 490 | 44 | 67 | 202 | 33 | 20 | 268 |
| 100% | 311 | 8 | 43 | 197 | 34 | 20 | 44 |
| **Total number DiMEs** | 4410 | 4805 | 535 | 1193 | 188 | 101 | 961 |
| **Total unique DiMEs** | 3547 | 3907 | 403 | 364 | 89 | 41 | 767 |
| **Total no. metabolites** | 50 | 29 | 50 | 52 | 40 | 40 | 47 |
| **Total no. substrates** | 41 | 25 | 44 | 46 | 30 | 32 | 39 |
| **Total no. products** | 29 | 19 | 12 | 14 | 12 | 9 | 21 |

Table S3: Added reactions to the models.

| Organism | Added reactions IDs | Added reactions formulas |
| --- | --- | --- |
| *S. alvi* | ASN2 | atp_c + nh4_c + asp__L_c *↔*h_c + amp_c + ppi_c + asn__L_c |
|  | PDX5PO2 | nad_c + pdx5p_c *↔* h_c + nadh_c + pydx5p_c |
|  | CA2abcpp | atp_c + h2o_c + ca2_p *→*adp_c + pi_c + h_c + ca2_c |
| *G. apicola* | 4HTHRA | 4hthr_c *↔*gly_c + gcald_c |
| *L. apis* | CU2tex | cu2_e *↔*cu2_p |
|  | DHBS | atp_c + h_c + 23dhb_c *↔*ppi_c + 23dhba_c |
|  | DHBSH | h2o_c + 23dhbzs_c *↔*ser__L_c + 23dhb_c |
|  | ENTCS | 3 23dhba_c + 3 seramp_c *↔*9 h_c + 6 amp_c + enter_c |
|  | ENTERES2 | 3 h2o_c + feenter_c *↔*3 h_c + fe3_c + 3 23dhbzs_c |
|  | SERASr | atp_c + h_c + ser__L_c *↔*ppi_c + seramp_c |
| *L. kullabergensis* | DHBS | atp_c + h_c + 23dhb_c *↔*ppi_c + 23dhba_c |
|  | DHBSH | h2o_c + 23dhbzs_c *↔*ser__L_c + 23dhb_c |
|  | ENTCS | 3 23dhba_c + 3 seramp_c *↔*9 h_c + 6 amp_c + enter_c |
|  | ENTERES2 | 3 h2o_c + feenter_c *↔*3 h_c + fe3_c + 3 23dhbzs_c |
|  | SERASr | atp_c + h_c + ser__L_c *↔*ppi_c + seramp_c |
|  | FRD2 | fum_c + mql8_c *↔*succ_c + mqn8_c |
|  | PDX5PO2 | nad_c + pdx5p_c *↔*h_c + nadh_c + pydx5p_c |
|  | MNt2pp | h_e + mn2_p *→*h_c + mn2_c |
|  | DHBS | atp_c + h_c + 23dhb_c *↔*ppi_c + 23dhba_c |
|  | MNt2pp | h_p + mn2_p *↔*h_c + mn2_c |
| *L. mellifer* | ZNabcpp | atp_c + h2o_c + zn2_p *↔*adp_c + h_c + pi_c + zn2_c |
| *B. asteroides* | INOSTO | h2o_c + h_c + glcur_c *↔*o2_c + inost_c |
| *R. ferrireducens* | PDX5PO2 | nad_c + pdx5p_c *↔*h_c + nadh_c + pydx5p_c |

Table S4: Statistics of the models used for the core honeybeegut microbiome studies with their thermodynamic coverage in % for metabolites and reactions in the model.

|  | | | thermodynamic coverage | |
| --- | --- | --- | --- | --- |
| model name | number metabolites | number reactions | % metabolites | % reactions |
| Bifidobacterium_asteroides_PRL2011 | 743 | 970 | 89% | 74% |
| Gilliamella_apicola_wkB1 | 1425 | 1994 | 90% | 60% |
| Lactobacillus_apis_Hma11 | 662 | 850 | 86% | 68% |
| Lactobacillus_kullabergensis_Biut2 | 789 | 997 | 79% | 59% |
| Lactobacillus_mellifer_Bin4 | 777 | 979 | 79% | 63% |
| Lactobacillus_mellis_Hon2 | 748 | 972 | 85% | 68% |
| Snodgrassella_alvi_wkB2 | 1015 | 1383 | 86% | 64% |

Table S5: Catabolite repression upper bound used for each model in the honeybee gut microbiome.

| model | $Cu,b$ |
| --- | --- |
| Bifidobacterium_asteroides_PRL2011 | 3 |
| Gilliamella_apicola_wkB1 | 3 |
| Lactobacillus_apis_Hma11 | 3 |
| Lactobacillus_kullabergensis_Biut2 | 3 |
| Lactobacillus_mellifer_Bin4 | 4 |
| Lactobacillus_mellis_Hon2 | 3 |
| Snodgrassella_alvi_wkB2 | 3 |
